# Ion transport peptide regulates body water balance via a receptor guanylyl cyclase in the *Drosophila* hindgut

**DOI:** 10.1101/2025.10.24.684309

**Authors:** Akira Watanabe, Takashi Koyama, Olga Kubrak, Michael James Texada, Megumi Furumitsu, Eiko Iwakoshi-Ukena, Roilea Maxson, Taishi Yoshii, Shu Kondo, Naoki Yamanaka, Kazuyoshi Ukena, Kim Rewitz, Ryusuke Niwa, Naoki Okamoto

**Affiliations:** Degree Programs in Life and Earth Sciences, Graduate School of Science and Technology, University of Tsukuba, Tennodai 1-1-1, Tsukuba, Ibaraki 305-8572, Japan; Department of Biology, Section for Cell and Neurobiology, University of Copenhagen, Universitetsparken 15, Building 3, 2100 Copenhagen, Denmark; Graduate School of Integrated Sciences for Life, Hiroshima University, Hiroshima 739-8521, Japan; Department of Entomology, Institute for Integrative Genome Biology, University of California, Riverside, 900 University Ave, Riverside, CA 92521, USA; Graduate School of Environmental, Life, Natural Science and Technology, Okayama University, Okayama, 700-8530, Japan; Department of Biological Science and Technology, Faculty of Advanced Engineering, Tokyo University of Science, 6-3-1 Niijuku, Katsushika-ku, Tokyo 125-8585, Japan; Life Science Center for Survival Dynamics, Tsukuba Advanced Research Alliance (TARA), University of Tsukuba, Tennodai 1-1-1, Tsukuba, Ibaraki 305-8577, Japan

## Abstract

Maintaining internal water balance, including ionic and osmotic balance, is critical for animal development and survival. In insects, internal water balance is regulated through the Malpighian tubules and the hindgut, which transport water and ions across their epithelia under the regulation of multiple peptide hormones. One of these, ion transport peptide (ITP), a member of the highly conserved crustacean hyperglycemic hormone superfamily among ecdysozoans, is an anti-diuretic hormone in insects, but its mechanism of action remains largely unclear. Here, we show that the short amidated isoform of ITP (saITP), secreted from brain neurosecretory cells, regulates water absorption via the receptor guanylyl cyclase (rGC) Gyc76C in the hindgut of the fruit fly *Drosophila melanogaster*. Both *ITP* and *Gyc76C* are evolutionarily conserved and are essential for larval survival in *D. melanogaster*, as mutation or knockdown of either gene results in early larval lethality. In *Drosophila* S2 cells, synthetic saITP increases intracellular cyclic GMP (cGMP) in a Gyc76C-dependent manner, an effect abolished by deletion of its putative ligand-binding domain. Using the fluorescent cGMP biosensor RedcGull in *ex vivo*-cultured hindguts, we found that saITP increases epithelial cGMP levels, and this response requires Gyc76C. Consistently, saITP promotes hindgut water reabsorption via Gyc76C. Thus, saITP acts via Gyc76C in a brain-hindgut neuroendocrine axis that regulates internal water balance during development. Our study provides new insight into the neuroendocrine control of osmoregulation in ecdysozoans and supports a broader role for rGCs in peptide-hormone signaling.

## Introduction

Maintaining water homeostasis, including the regulation of ionic and osmotic balance, is essential for the development and survival of all animals (1). Although the underlying mechanisms vary depending on their habitat, many animals, including insects, rely on endocrine systems to coordinate systemic water and ion balance (2, 3). In insects, water and ion balance is primarily maintained by the Malpighian tubules and the hindgut, organs functionally analogous to the vertebrate kidney and large intestine. These organs, like their mammalian counterparts, dynamically adjust water and ion reabsorption or excretion in response to multiple neuropeptides and peptide hormones (3–5).

Ion transport peptide (ITP), a member of the crustacean hyperglycemic hormone (CHH) superfamily, is important for water homeostasis and is widely conserved among ecdysozoans (6, 7). ITP was initially identified in the desert locust *Schistocerca gregaria* as a hormone promoting ion and fluid transport across the hindgut epithelium (8–10). Subsequent studies established ITP as an anti-diuretic hormone in the fruit fly *Drosophila melanogaster*, particularly under conditions of dehydration or desiccation stress (11). In addition to its role in water homeostasis, ITP exhibits significant pleiotropy in insects, regulating metabolism, molting, circadian rhythm, and reproduction (12–20). This functional diversity may arise from the alternative splicing of the *ITP* gene, which leads to the translation of multiple isoforms. At least two evolutionarily conserved isoforms are produced from the *ITP* gene: a short, C-terminally amidated isoform (hereafter referred to as short amidated ITP or saITP) and a longer, non-amidated isoform (ITP-like or ITPL) (6). In *S. gregaria*, only saITP was found to promote water transport in the hindgut (21). Similar findings were recently reported in the yellow fever mosquito *Aedes aegypti*, in which knockdown of *saITP* but not *ITPL* transcripts resulted in abnormal water homeostasis (19). Although these results lend further support to a conserved role for saITP in ion and fluid transport across insect species, the molecular mechanisms that underlie the differential actions of the saITP and ITPL isoforms remain unclear, especially the relevant receptor(s) mediating their actions. Some studies have suggested that CHH superfamily members act via receptor guanylyl cyclases (rGCs), because cyclic GMP (cGMP) is the predominant second messenger in osmoregulation (22, 23), whereas other studies have suggested G protein-coupled receptors (GPCRs) as the transducers of CHH-family peptide activity (24). In the silkmoth *Bombyx mori*, three GPCRs reportedly respond to saITP and ITPL in cell culture (25, 26). More recently, another group found that *Drosophila* ITPL2 exerts an anti-diuretic effect in the Malpighian tubules via the tachykinin-related peptide (TRP) receptor TkR99D (17). The phylogenetic distribution of these GPCRs do not fully match the evolutionary conservation of the CHH superfamily. Moreover, the TRP receptor exhibits a higher affinity for its canonical ligand, TRP, than for ITPL (26). This suggests the existence of one or more as-yet-unidentified, *bona fide* ITP receptors.

In this study, we used the *D. melanogaster* model to identify an evolutionarily conserved receptor for saITP, the ITP isoforms previously implicated in regulating water homeostasis (19, 21). We demonstrate that saITP, secreted from neurosecretory cells (NSCs) in the brain, regulates water homeostasis through the rGC Gyc76C in the hindgut of *D. melanogaster*. Both *ITP* and *Gyc76C* are essential for larval survival and are evolutionarily conserved among ecdysozoans. In *Drosophila* S2 cells and *ex vivo* cultured hindguts, we found that application of synthetic saITP induced a robust increase in intracellular cGMP through Gyc76C and that this increase required its putative ligand-binding domain (LBD). We also found that hindgut-specific *Gyc76C* knockdown eliminated the water reabsorption effect of saITP. Thus, our results indicate that saITP acts via Gyc76C in a neuroendocrine circuit operating through a brain–hindgut axis to maintain water balance during development.

## Results

### saITP is a neuroendocrine peptide specifically produced in the brain

In many insects, saITP is produced primarily by NSCs in the brain, whereas *ITPL* is additionally expressed in diverse peripheral tissues (6, 12, 15, 18, 19, 25, 27). In *D. melanogaster*, the *ITP* gene encodes, through alternative splicing, a single saITP and two ITPLs, ITPL1 and ITPL2 (27). In a tissue-specific larval expression analysis, we found that *ITPL1* and *ITPL2* are broadly expressed in peripheral tissues (Fig. S1A), whereas *saITP* expression is restricted to the larval central nervous system (CNS) during development (Fig. 1A and S1A). To characterize the *saITP*-expressing cells of the CNS, we generated a *saITP::T2A::GAL4* driver line that mimics endogenous *saITP* expression. Consistent with previous reports (27), we found evidence of *saITP* expression in several NSC populations, including four pairs of ITP-immunoreactive protocerebral (*ipc-1*) neurons in the larval brain, a single pair of ITP-immunoreactive subesophageal (*isog*) neurons in the subesophageal zone, and a few pairs of ITP-immunoreactive abdominal ganglion (*iag*) neurons in the ventral nerve cord (Fig. 1B). Notably, the axons of the *ipc-1* neurons project to the neurohemal organ called the *corpora cardiaca* (CC), which may be a site of saITP secretion into the hemolymph. Indeed, we confirmed via immunostaining with an anti-saITP antibody that saITP is strongly localized within the CC-targeting projections of the *ipc-1* neurons (Fig. 1C). These results suggest that, during larval life, saITP is predominantly produced by NSCs in the CNS and released into circulation at sites near the CC. This is consistent with findings in *S. gregaria*, in which hindgut ion-transport activity is stimulated specifically by extracts from the brain and CC (8–10).

**Figure 1:**
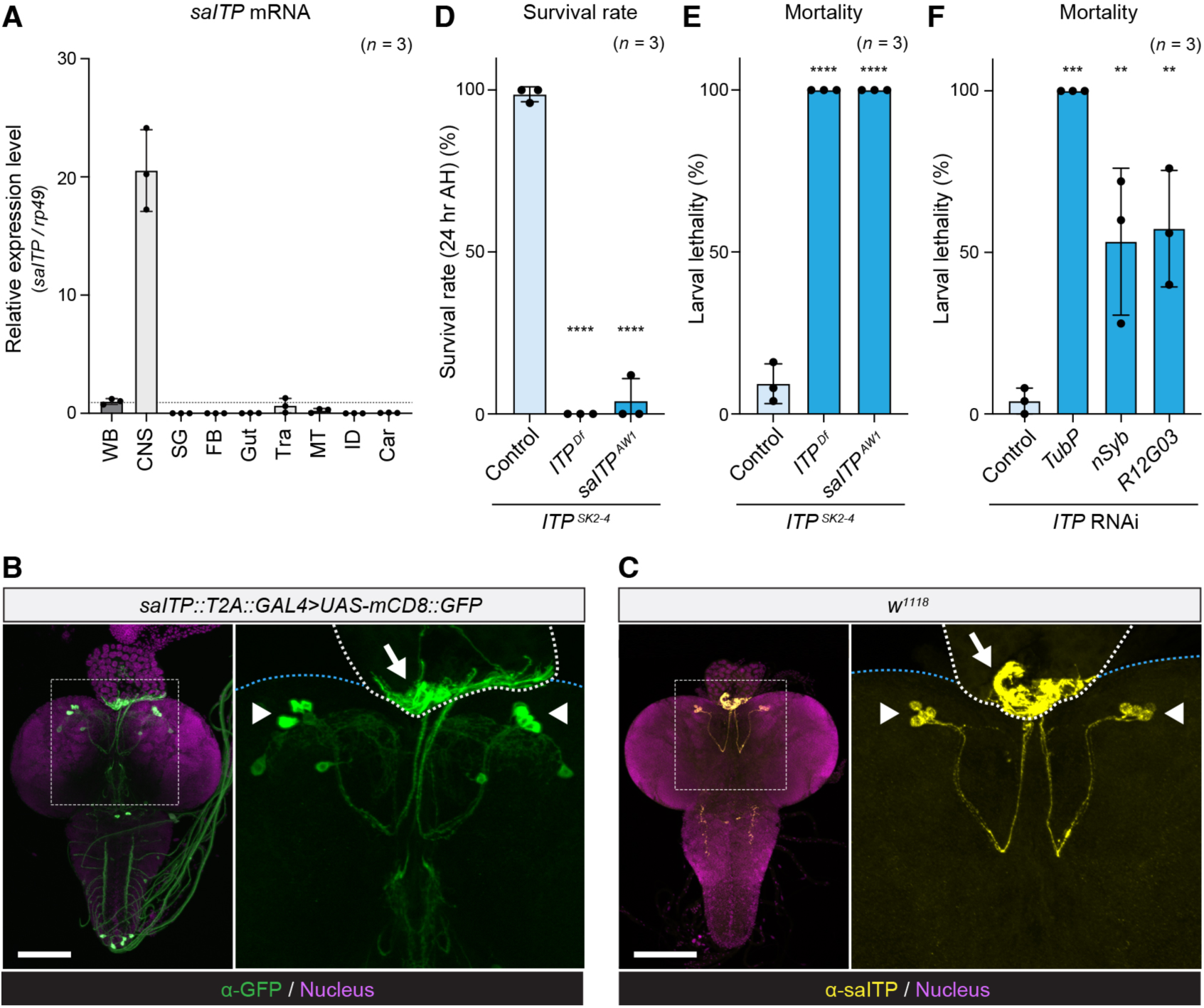
saITP produced by the CNS is essential for survival during development. **A-C**, saITP is specifically produced by brain NSCs. **A**, Relative expression levels of *saITP* in various tissues, as assessed by RT-qPCR. WB, whole body; CNS, central nervous system; SG, salivary glands; FB, fat body; Tra, tracheae; MT, Malpighian tubules; ID, imaginal discs; Car, carcass. Values are normalized to expression in WB. **B** and **C**, *saITP::T2A::GAL4* expression (green) (**B**) and saITP immunoreactivity (yellow) (**C**) in the dissected CNS of wandering third instar larvae. Nuclei were stained with DAPI (magenta). *saITP::T2A::GAL4* was visualized by *UAS-mCD8::GFP* (**B**). The right image in each panel shows a magnified view of the area outlined by the dotted square. The arrowheads indicate the *ipc-1* neurons, and the arrow indicates projections to the CC. Scale bars, 100 µm. **D** and **E**, Survival rate 24 hrs of hatching (AH) (**D**) and larval mortality rates (**E**) of control (*ITP^SK2-4^*/*+*), *ITP* mutant (*ITP^SK2-4^/ITP^Df^*), and *saITP* isoform-specific mutant (*ITP^SK2-4^/saITP^AW1^*) larvae. **F**, Mortality rates of larvae with tissue-specific *ITP* knockdown using the following *GAL4* driver lines: *TubP* (ubiquitous), *nSyb* (pan-neuronal), and *R12G03* (*ipc-1* neurons). Each *GAL4* line was crossed to *UAS-ITP RNAi*. Larvae were reared at 25 individuals per vial, and the data represent the results of three independent experiments (**D**, **E**, and **F**). Data are presented as means ± SD. Sample sizes (*n*) are indicated on each graph. \*\**p* < 0.01, \*\*\**p* < 0.001, \*\*\*\**p* < 0.0001, as assessed by one-way ANOVA with Dunnett’s test for multiple comparisons to the control (**D**, **E**, and **F**).

### Brain-derived saITP is essential during development

Given the critical role of water homeostasis in animal development and viability, we hypothesized that *ITP* loss-of-function during *Drosophila* development would likely induce lethality. Indeed, both a CRISPR-generated mutant lacking all *ITP* isoforms (*ITP^SK2-4^*) and ubiquitous *ITP* knockdown caused lethality during the first larval instar within 24 hrs of hatching (Fig. 1D), with no larvae able to progress to the pupal stage (Fig. 1E and F). This is consistent with a previous report that ubiquitous knockdown of *ITP* causes developmental lethality (16). Furthermore, both pan-neuronal (*nSyb-GAL4*) and *ipc-1* neuron-specific (*R12G03-GAL4*) knockdown of *ITP* led to larval lethality (Fig. 1E and S1B). To determine whether this phenotype was caused by loss of *saITP*, rather than the other variants, we used CRISPR/Cas9 to generate a *saITP* isoform-specific mutant. As with the *ITP* null mutant, the resulting *saITP* mutant (*saITP^AW1^*) exhibited strong lethality during the first larval instar (Fig. 1D and E). Together, these results suggest that brain-derived saITP is essential for survival during larval development.

### Identification of evolutionarily conserved candidate receptors for saITP

Previous studies in *B. mori* identified two GPCRs, *Bombyx* orphan neuropeptide GPCR (BNGR)-A2 and BNGR-A34, as candidate saITP receptors (25). *BNGR-A2* is conserved only within certain Lepidopteran species, whereas *BNGR-A34* orthologs are present in a number of dipteran and coleopteran genomes, including that of *D. melanogaster* (25). The phylogenetic distribution of these GPCRs, however, does not match the broad evolutionary conservation of the CHH superfamily across arthropods (25). In addition, BNGR-A24 and TkR99D, which are potential receptors for ITPL in *B. mori* and *D. melanogaster*, respectively (17, 25), encode TRP receptors (26, 28) with higher affinity for TRP ligands than for ITPL (26). To determine whether these GPCRs are *bona fide* saITP receptors in *D. melanogaster*, we analyzed the developmental phenotypes of mutants lacking *CG30340* (the *Drosophila* ortholog of *BNGR-A34*) and *TkR99D*. If either of these GPCRs is an saITP receptor, loss of its function should phenocopy the loss of saITP and drive larval lethality. Both *CG30340^attP^* and *TkR99D^attP^* mutants, however, were viable and without any evidence of larval lethality (Fig. S2A).

Recognition of peptide hormones by their cognate receptors often relies on specific structural motifs dictated by conserved amino acids (29–31). As a member of the CHH superfamily, saITP contains six conserved cysteine residues that likely contribute to forming a specific three-dimensional conformation via disulfide bridges (Fig. 2A) (6, 32, 33). To identify candidate receptors that recognize this structural motif, we used the ScanProsite tool (https://prosite.expasy.org/scanprosite/) to search for neuropeptides and peptide hormones in the *Drosophila* genome with a cysteine pattern similar to that of saITP. This survey revealed a partial match between saITP and eclosion hormone (EH), another peptide hormone that contains six conserved cysteines (Fig. 2A). Notably, both peptides share an identical arrangement of four internal cysteine residues (C-X_2_-C-X_12_-C-X_3_-C) (Fig. 2A), suggesting a similar tertiary structure and possibly recognition by the same class of receptors. Since EH acts via an rGC in *S. gregaria*, *D. melanogaster*, and the oriental fruit fly *Bactrocera dorsalis*, (34–36), we hypothesized that saITP may also signal via an rGC.

**Figure 2:**
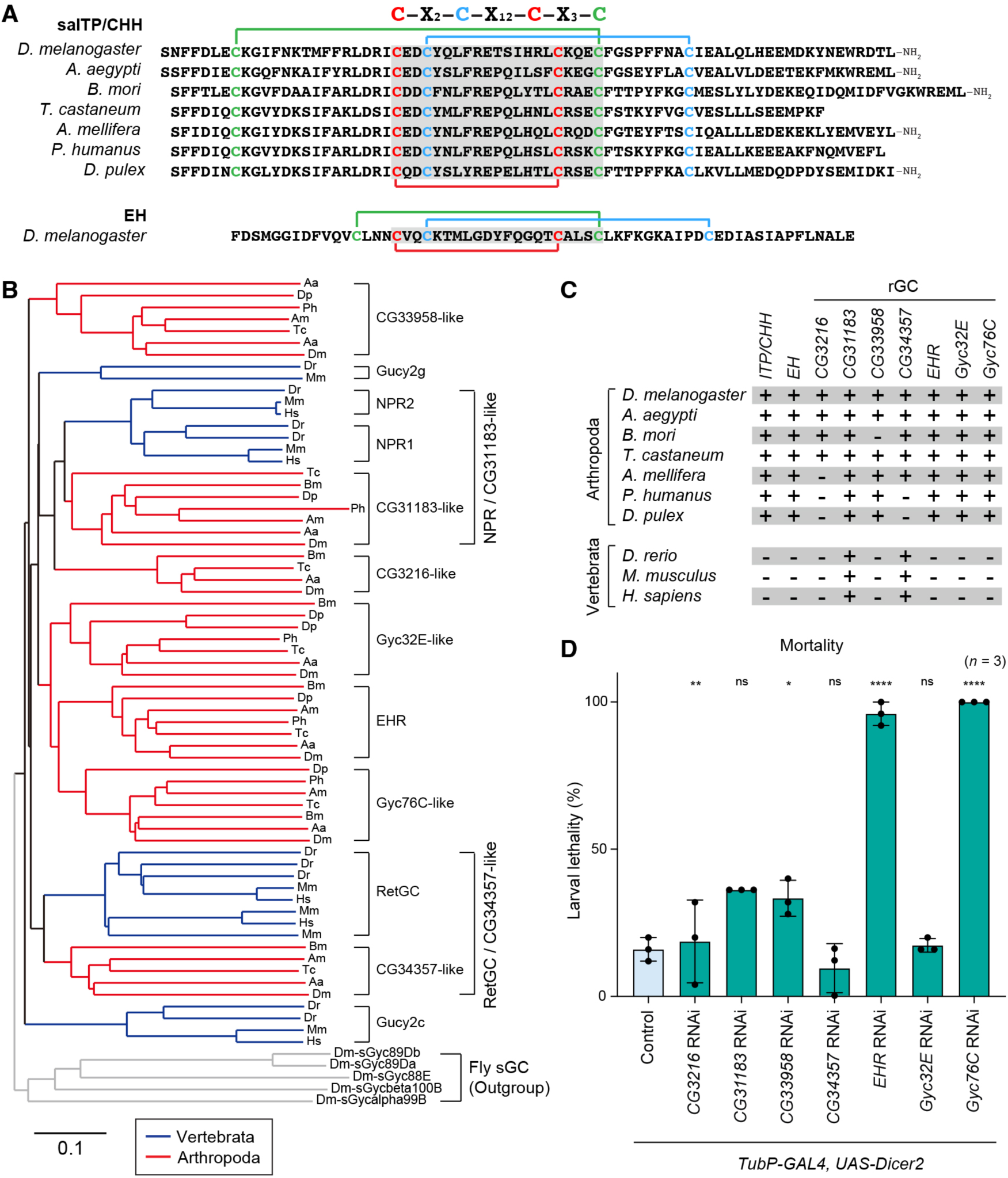
Phylogenetic analysis and *in vivo* RNAi screening to identify candidate saITP receptors. **A**, Multiple alignment of the amino-acid sequences of mature saITP/CHH peptides in holometabolous insects (*D. melanogaster*; yellow fever mosquito, *Aedes aegypti*; silk moth, *Bombyx mori*; red flour beetle, *Tribolium castaneum*; western honey bee, *Apis mellifera*), a hemimetabolous insect (human body louse, *Pediculus humanus*), and a crustacean (water flea, *Daphnia pulex*), and EH from *D. melanogaster*. Peptides encoded by all *ITP* orthologs contain six highly conserved cysteine residues that form three disulfide bonds. The shaded regions in the amino-acid sequences indicate the identical arrangement of four internal cysteine residues (C-X_2_-C-X_12_-C-X_3_-C) in saITP/CHH and EH. **B** and **C**, Phylogenetic analysis of rGC proteins in the genomes of arthropods and vertebrates. **B**, Neighbor-joining tree constructed using the full amino-acid sequences of rGC proteins from vertebrates (human, *Homo sapiens*; mouse, *Mus musculus*; zebrafish, *Danio rerio*) and arthropods (*D. melanogaster*, *A. aegypti*, *B. mori*, *T. castaneum*, *A. mellifera*, *P. humanus*, and *D. pulex*). Protein names and GenBank accession numbers are listed in Table S1. The scale bar represents an evolutionary distance of 0.1 amino-acid substitutions per site. **C**, Presence or absence of ITP/CHH, EH, and rGCs in selected arthropods and vertebrate species. Orthologs in the indicated species were identified by BLAST analysis of their genome assemblies and are indicated by “+” (presence) or “−” (absence). **D**, Larval mortality rates following ubiquitous knockdown of genes encoding rGCs. Larvae were reared at 25 individuals per vial, and the data represent the results of three independent experiments. *TubP-GAL4>UAS-Dicer-2* was used as a ubiquitously expressing driver. Data are presented as means ± SD. Sample sizes (*n*) are shown in each graph. \**p* < 0.05, \*\**p* < 0.01, \*\*\*\**p* < 0.0001, as assessed by one-way ANOVA with Dunnett’s test for multiple comparisons to the control (**D**). ns: not significant (*p* > 0.05).

The *Drosophila* genome encodes seven rGCs: *CG3216, CG10738* (hereafter referred to as *EH receptor* or *EHR*), *CG31183*, *CG33958*, *CG34357*, *Gyc32E*, and *Gyc76C*. EHR functions as a receptor for EH (35), but the other receptors’ cognate ligands remain unknown. To assess the evolutionary conservation of these receptors, we examined the orthologs of these seven rGC-encoding genes across a broad range of arthropods and vertebrates. Four of the rGC-encoding genes—*CG31183*, *EHR*, *Gyc32E*, and *Gyc76C*—are highly conserved across all examined arthropod species (Fig. 2C). Notably, orthologs of three rGCs, such as *EHR*, *Gyc32E*, and *Gyc76C*, are absent in vertebrates but are uniformly present within Arthropoda, paralleling the phylogenetic distribution of the CHH superfamily (Fig. 2C).

To assess the potential for each of the seven *Drosophila* rGCs to function as a receptor for saITP, we ubiquitously knocked down all rGCs in the larval stage and analyzed lethality as a readout. Knockdown of *EHR* or *Gyc76C* resulted in near-total larval lethality (Fig. 2D), an outcome that was confirmed using an alternative RNAi line for each gene (Fig. S2B). Given that EH and EHR are required for molting during larval development in *D. melanogaster* (35, 37, 38), *EHR* knockdown-induced lethality is consistent with its known function. Thus, Gyc76C emerged as the top saITP-receptor candidate.

### Synthetic saITP elevates intracellular cGMP levels via Gyc76C

To determine whether saITP activates Gyc76C, we conducted *in vitro* assays in which chemically synthesized *Drosophila* saITP was applied to *Drosophila* S2 cells. This saITP contained the three disulfide bonds and C-terminal amidation that characterize active CHH-superfamily peptides (39, 40). Since members of the CHH superfamily, including saITP, signal via the second messenger cGMP (7, 41), we assessed receptor activation by measuring intracellular cGMP levels.

Addition of saITP to untransformed S2 cells did not induce a cGMP response (Fig. 3A). Overexpression of *Gyc76C* in S2 cells led to elevated basal cGMP levels, and the addition of synthetic saITP further increased cGMP levels significantly (Fig. 3A), indicating that exogenous Gyc76C is sufficient for saITP responses in these cells. In contrast, S2 cells overexpressing *EHR* did not respond to saITP (Fig. 3A). When we added an alternative reported ligand of Gyc76C, the VQQ peptide of Neuropeptide-like precursor 1 (Nplp1-VQQ) (42), we did not observe any increase in cGMP levels (Fig. S3A). This lack of second-messenger response suggests that Nplp1-VQQ does not activate Gyc76C under our assay conditions.

**Figure 3:**
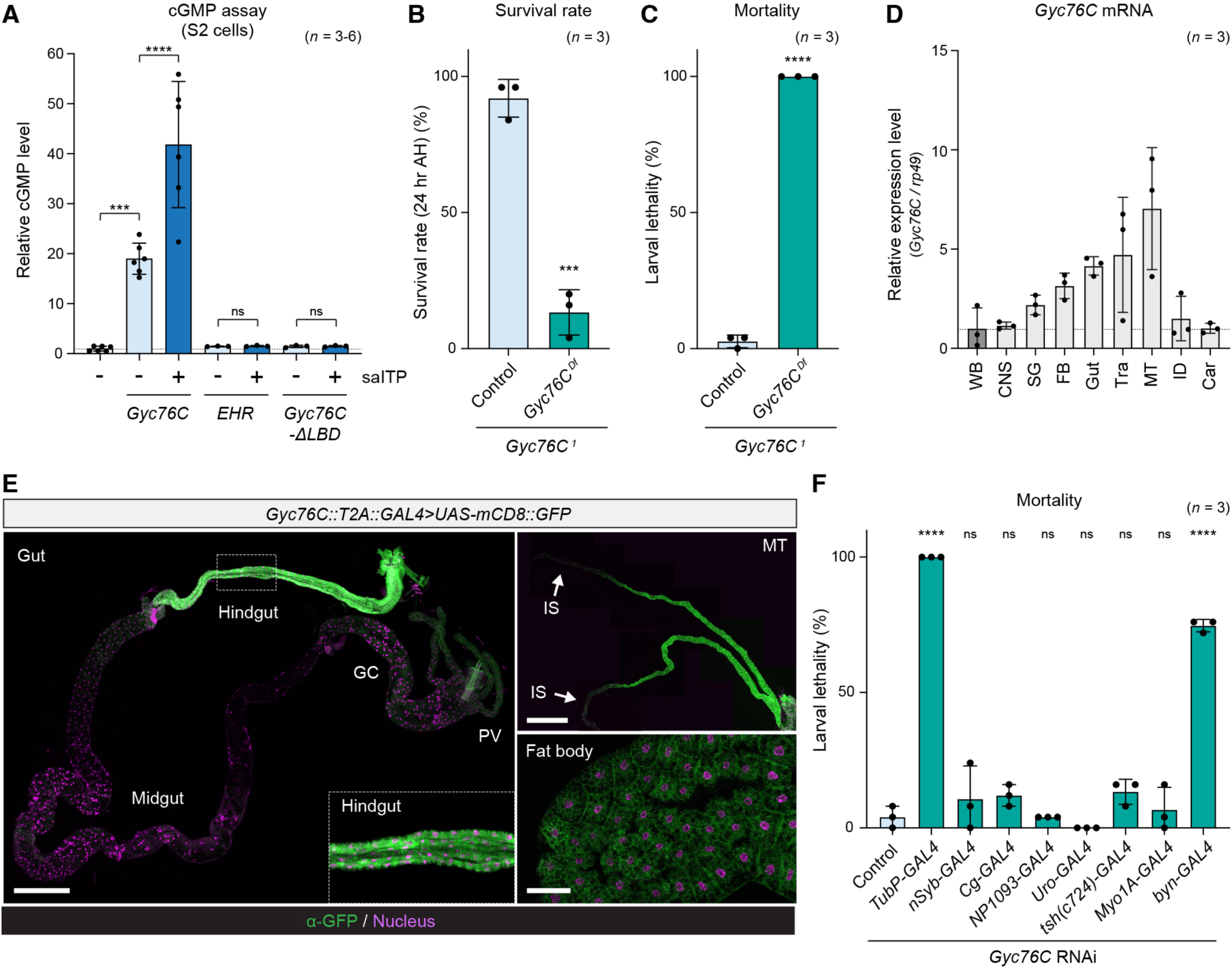
Gyc76C function in the hindgut is essential for survival during development. **A,** Intracellular cGMP levels in S2 cells expressing *Gyc76C*, *EHR*, and *Gyc76C-ΔLBD*, with (+) or without (−) the addition of synthetic saITP. *pActin-GAL4* was used to express each rGC in S2 cells. Values are shown relative to the control (−). **B** and **C**, Survival rate 24 hrs AH (**B**) and larval mortalities (**C**) for controls (*Gyc76C^1^*/*+*) and *Gyc76C* mutants (*Gyc76C^1^*/*Gyc76C^Df^*). **D**, Relative expression levels of *Gyc76C* in different tissues, as assessed by RT-qPCR. **E**, *Gyc76C::T2A::GAL4* expression patterns in different tissues of wandering third instar larvae. Nuclei were stained with DAPI (magenta). *Gyc76C::T2A::GAL4* was crossed to *UAS-mCD8::GFP*. Strong reporter expression was apparent in the hindgut, but only low levels were observed in the proventriculus (PV), gastric caeca (GC), and midgut. Reporter was also expressed in the Malpighian tubules (MT), expect for their initial segments (IS), and in the fat body. Scale bars, 500 µm (gut and MT) and 100 µm (fat body). **F**, Larval mortality rates following tissue-specific knockdown of *Gyc76C* driven by the following *GAL4* driver lines: *TubP-GAL4* (ubiquitous), *nSyb-GAL4* (pan-neuronal), *Cg-GAL4* (fat body), *NP1093-GAL4* (principal cells of the Malpighian tubules), *Uro-GAL4* (principal cells of the Malpighian tubules in the main segments), *tsh(c724)-GAL4* (stellate cells of the Malpighian tubules), *Myo1A-GAL4* (gut enterocytes), and *byn-GAL4* (hindgut). Each *GAL4* line was crossed to *UAS-Gyc76C RNAi*. Larvae were reared at 25 individuals per vial, and the data represent the results of three independent experiments (**B**, **C** and **F**). Data are presented as means ± SD. Sample sizes (*n*) are shown in each graph. \*\*\**p* < 0.001, \*\*\*\**p* < 0.0001, as assessed by one-way ANOVA with Tukey’s test for multiple comparisons (**A**), unpaired *t* tests (two-tailed) (**B** and **C**), or one-way ANOVA with Dunnett’s test for multiple comparisons to the control (**F**). ns: not significant (*p* > 0.05).

To confirm whether the cGMP response to saITP observed in *Gyc76C*-expressing cells requires direct binding to Gyc76C, we generated a deletion construct lacking the predicted extracellular ligand binding domain (LBD) of Gyc76C (*Gyc76C-ΔLBD*). S2 cells expressing this construct *Gyc76C-ΔLBD* showed neither the basal cGMP elevation nor the saITP-induced cGMP response that we observed in controls (Fig. 3A), indicating that the LBD is required for Gyc76C-mediated saITP response and suggesting that saITP exerts its effect via a direct interaction with the Gyc76C LBD.

The increase in basal cGMP levels in S2 cells overexpressing *Gyc76C* even before saITP addition suggests that S2 cells may secrete endogenous ligands that activate Gyc76C in an autocrine or paracrine manner. To test this, we knocked down endogenous *ITP* in *Gyc76C*-expressing cells and observed a significant reduction in basal cGMP concentration (Fig. S3B), suggesting that S2 cells produce and secrete ITP peptides capable of activating Gyc76C in S2 cells. Indeed, the FlyAtlas expression database (http://flyatlas.org/atlas.cgi) reports S2-cell expression of *ITP*.

To further evaluate whether Gyc76C alone is sufficient for saITP signal transduction, we performed ligand-stimulation assays using human HEK293T cells overexpressing *Drosophila Gyc76C*. In these cells, however, saITP application did not induce a significant increase in cGMP levels (Fig. S3C). This lack of effect suggests that saITP signaling via Gyc76C requires additional co-factors or cell-type-specific machinery that is present in S2 cells but absent in HEK293T cells.

### Gyc76C function in the hindgut is essential for survival during development

Since we found that ubiquitous knockdown of *Gyc76C* phenocopied the lethality of *saITP* mutants (Fig. 2D and S2B), we next generated a *Gyc76C* loss-of-function mutant using CRISPR/Cas9.

Consistent with our RNAi results, we found that *Gyc76C* mutant (*Gyc76C^1^*/*Df*) did not survive beyond 24 hrs post-hatching (Fig. 3B), with no larvae progressing to puparium formation (Fig. 3C). This developmental phenotype closely resembles that of *saITP* mutants.

To identify where this lethality phenotype might arise, we performed a tissue-specific analysis of *Gyc76C* expression. We found broad expression of *Gyc76C* in the CNS and peripheral tissues (Fig. 3D). Using a *Gyc76C::T2A::GAL4* knock-in line, which mimics endogenous *Gyc76C* expression, we observed reporter expression in multiple tissues, including the gut, Malpighian tubules, and fat body (Fig. 3E). Within the gut, *Gyc76C* showed particularly high expression in the hindgut (Fig. 3E), which is a known target tissue of saITP in the desert locust (33). We next assessed the developmental viability of strains carrying *GAL4* transgenes driving targeted in lines with tissue-specific *Gyc76C* knockdown of *Gyc76C* in relevant tissues. Strikingly, we found that hindgut-specific knockdown of *Gyc76C* induced significant larval lethality (Fig. 3F). In contrast, *Gyc76C* knockdown in other osmoregulatory tissues, including the principal cells (*NP1093-GAL4* and *Uro-GAL4*) and stellate cells (*tsh[c724]-GAL4*) of the Malpighian tubules, had no effect on developmental viability (Fig. 3F). These findings demonstrate that Gyc76C in the hindgut is essential for survival during development and suggest that the hindgut as the primary target of saITP in *D. melanogaster*.

### saITP acts directly in the hindgut epithelium through Gyc76C

To directly determine whether saITP acts on the hindgut, we performed *ex vivo* live imaging of dissected hindguts after saITP addition. To visualize changes in intracellular cGMP levels, we established a *UAS* transgenic line to permit tissue-specific expression of the red fluorescent cGMP sensor RedcGull (43). When we added synthetic saITP to third instar larval hindguts expressing *RedcGull*, we observed a robust increase in cGMP signal in hindgut epithelial cells (Fig. 4A).

**Figure 4:**
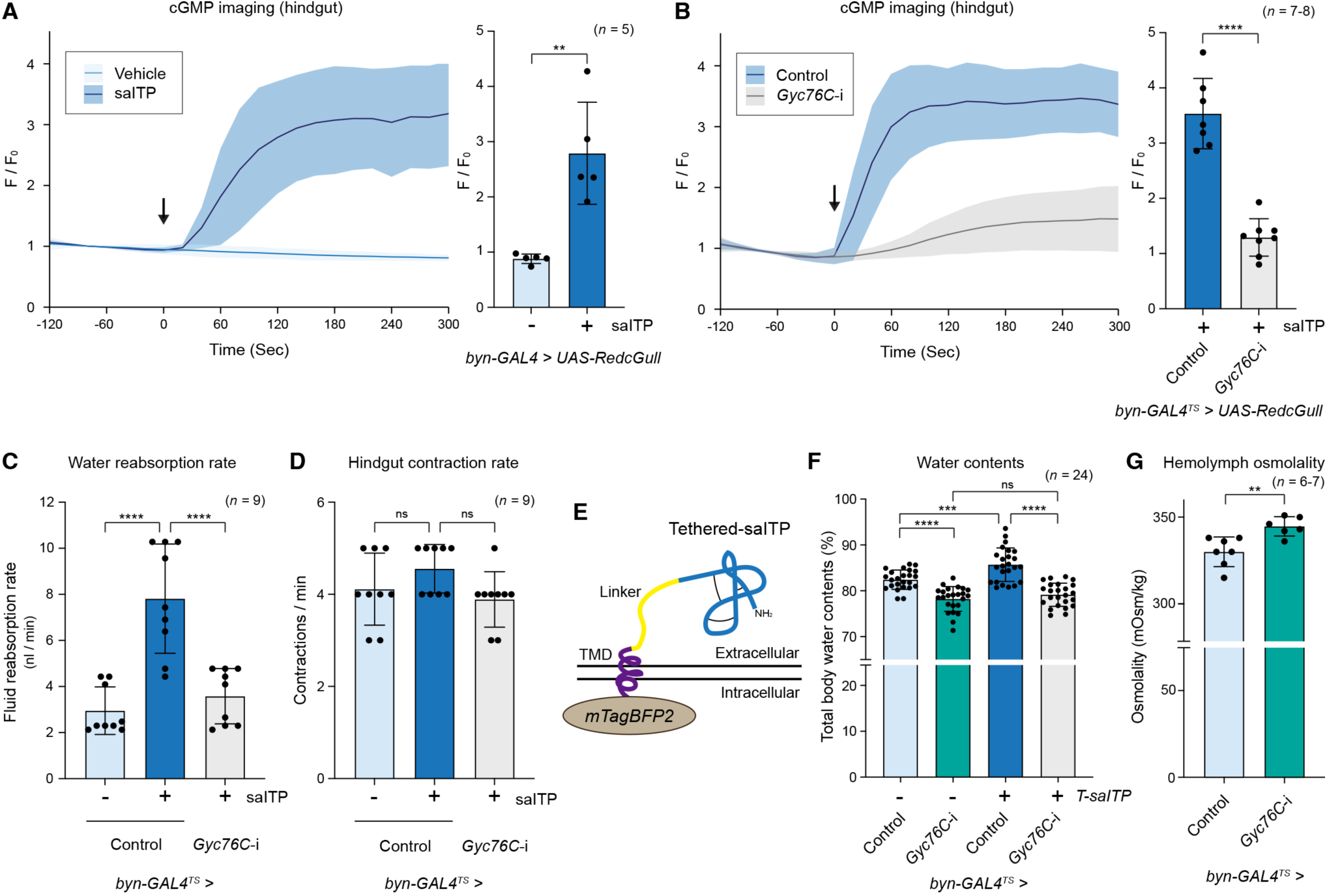
saITP acts directly on the hindgut epithelium to promote water reabsorption via Gyc76C. **A** and **B,** saITP acts directly on the hindgut to increase intracellular cGMP levels via Gyc76C. **A**, RedcGull fluorescence intensity in the hindgut of larvae with (+) or without (−) addition of synthetic saITP. **B**, RedcGull fluorescence intensity in the hindgut of control and *Gyc76C* RNAi (*Gyc76C*-i) larvae treated with saITP. Left graphs show temporal changes in RedcGull fluorescence intensity. Right graphs show quantification of RedcGull fluorescence intensity at 120 seconds post-stimulation. The average fluorescence intensity immediately before the addition of the reagent was defined as F_0_ and used to normalize the values at all other time points. *byn-GAL4* (**A**) and *byn-GAL4^TS^*(**B**) were used as hindgut-specific drivers. **C** and **D**, water reabsorption rates (**C**) and contraction rates (**D**) in the hindgut of control and *Gyc76C*-i larvae with (+) or without (−) addition of synthetic saITP. **E**, Schematic structure of the engineered membrane-tethered saITP protein. From the N terminus, the construct comprised mTagBFP2, a transmembrane domain, and saITP connected via a linker sequence, producing a membrane-anchored version of saITP that acts on its receptor in a constitutive autocrine (or potentially juxtacrine) manner. **F**, Total body water content in control and *Gyc76C-i* wandering third instar larvae with (+) or without (−) expression of *tethered-saITP* (*T-saITP*). **G**, Hemolymph osmolality (mOsm/kg) in control and *Gyc76C-i* third instar larvae. In all *Gyc76C* knockdown experiments, larvae were reared at 18 °C during the early larval stages (L1 and L2) and shifted to 29 °C from 0 hr after third instar ecdysis to induce RNAi from the third instar. Data are presented as means ± SD. Sample sizes (*n*) are shown in each graph. \*\**p* < 0.01, \*\*\**p* < 0.001, \*\*\*\**p* < 0.0001, as assessed by unpaired *t* tests (two-tailed) (**A**, **B** and **G**) and one-way ANOVA with Tukey’s test for multiple comparisons (**C, D** and **F**). ns: not significant (*p* > 0.05).

Notably, this response was strongly attenuated by simultaneous hindgut-specific knockdown of *Gyc76C* (Fig. 4B). These results demonstrate that saITP acts directly on the hindgut to increase intracellular cGMP levels in a Gyc76C-dependent manner.

To determine whether other *Gyc76C*-expressing tissues also respond to saITP, we next expressed *RedcGull* in the fat body, which expresses high levels of *Gyc76C* (Fig. 3D and E). We then monitored cGMP levels after saITP treatment. In contrast to the strong response induced in the hindgut, saITP application to the fat body did not induce changes in cGMP levels (Fig. S4).

Combined with the lack of observable saITP response in mammalian HEK293T cells expressing *Gyc76C* (Fig. S3C), these results suggest that Gyc76C alone is insufficient for saITP signaling. Instead, saITP appears to require tissue-specific co-receptors or cofactors present in the hindgut but not in the fat body.

### saITP promotes water reabsorption in the hindgut via Gyc76C

In insects, the hindgut is a key organ in the regulation of internal water balance as it performs the essential function of reabsorbing water and ions from excreta (3). Given the lethal phenotypes we observed with loss-of-function of either *saITP* or *Gyc76C*, we hypothesized that saITP-Gyc76C signaling promotes water reabsorption in the hindgut. To test this, we used an *ex vivo* water-reabsorption assay to quantify the rate of water uptake from the hindgut lumen (44, 45). We found that synthetic saITP significantly increased water reabsorption via the larval hindgut (Fig. 4C), paralleling the induction of cGMP signaling (Fig. 4B). Furthermore, this effect was abolished by hindgut-specific knockdown of *Gyc76C* (Fig. 4C). These effects strongly suggest that saITP acts in the hindgut via Gyc76C to promote water reabsorption. Notably, saITP had no effect on hindgut peristalsis (Fig. 4D), suggesting that saITP-mediated regulation of water reabsorption is independent of gut motility.

To further investigate whether saITP-Gyc76C signaling in the hindgut contributes to systemic water balance, we measured the total water content of third instar larvae. Since *Gyc76C* is expressed across multiple tissues, and endogenous saITP is a secreted factor that presumably can reach these locations, we could not use *saITP* overexpression to analyze any tissue-specific saITP actions. To overcome this, we engineered a membrane-tethered form of saITP (*UAS-tethered-saITP* or *T-saITP*). The membrane-tethered saITP would be anticipated to activate its receptor in a constitutive autocrine (or potentially juxtacrine) manner (Fig. 4E), without directly perturbing signaling in non-targeted tissues.

Knockdown of *Gyc76C* in the hindgut reduced larval water content compared to control animals (Fig. 4F), consistent with the results of the reabsorption experiment (Fig. 4C), suggesting that endogenous saITP-mediated Gyc76C signaling in the larval hindgut contributes to basal water homeostasis. On the other hand, targeted expression of *T-saITP* in the hindgut significantly increased larval water content, an effect consistent with enhanced water reabsorption in the hindgut epithelium (Fig. 4F). When we combined *T-saITP* expression with hindgut-specific knockdown of *Gyc76C*, the resulting larvae showed water content similar to that of animals expressing hindgut-specific *Gyc76C* knockdown alone (Fig. 4F), confirming that saITP regulates systemic water balance through hindgut Gyc76C. To further investigate whether Gyc76C in the hindgut contributes to systemic water balance, we measured larval hemolymph osmolality. Hindgut-specific *Gyc76C* knockdown increased osmolality relative to controls (Fig. 4G), consistent with reduced hindgut water reabsorption and therefore increased systemic dehydration. Interestingly, we also found that hindgut-specific expression of *T-saITP* severely impaired larval growth, resulting in smaller pupae (Fig. S5).

Although the mechanisms by which perturbation of systemic water balance might reduce body size remain unclear, hindgut-specific knockdown of *Gyc76C* completely suppressed this phenotype (Fig. S5), further supporting the notion that saITP acts on the hindgut through Gyc76C. Together, these findings demonstrate that saITP acts through Gyc76C to promote water reabsorption in the hindgut, thereby contributing to the maintenance of systemic water balance.

## Discussion

Terrestrial animals, including insects, must maintain proper internal water balance for both development and survival (3–5). In insects, water and ion homeostasis are primarily regulated by the Malpighian tubules and the hindgut, which dynamically adjust the reabsorption and excretion of water and ions in response to multiple neuropeptides and peptide hormones (3–5). However, our current understanding of water and ion homeostasis in insects, including *D. melanogaster*, has largely been derived from endocrine studies focusing on the Malpighian tubules (3–5). In contrast, the molecular mechanisms underlying water balance regulation in the hindgut remain poorly understood. In this study, we identified the saITP as a key neuroendocrine factor that acts on the hindgut to regulate systemic water balance in *D. melanogaster*. *saITP* is expressed in NSCs in the brain, and its expression is essential for larval survival. Furthermore, we demonstrated that saITP promotes cGMP production in the hindgut epithelium through activation of an rGC Gyc76C, thereby inducing water reabsorption. saITP was originally discovered as a bioactive peptide extracted and purified from the brain and CC of locusts that promotes hindgut water reabsorption (7). Our findings provide clear genetic evidence that, as it does in locusts, *Drosophila* saITP functions as a neuroendocrine factor acting on the larval hindgut to regulate systemic water balance. Our study reveals a critical brain–hindgut axis required for larval survival and provides new insights into the neuroendocrine control of water homeostasis during insect development.

Through phylogenetic and functional analyses, we identified Gyc76C as a candidate receptor for saITP in *D. melanogaster*. Another recent study independently implicated Gyc76C in saITP signaling in adult *D. melanogaster* by showing that saITP inhibition of LK-and DH31-stimulated Malpighian tubule fluid secretion depends on Gyc76C (46), supporting our findings. Using tissue-specific knockdown, we demonstrated that *Gyc76C* is essential in the hindgut for survival during development, but dispensable in other larval osmoregulatory tissues, such as the Malpighian tubules. Furthermore, using *ex vivo* live cGMP imaging and water-reabsorption assays, we demonstrated that saITP promotes water uptake in the hindgut epithelium via Gyc76C. Notably, we used a membrane-tethered saITP with spatially restricted activity to confirm its direct role in modulating water balance via Gyc76C in the hindgut epithelium. We also showed that the predicted extracellular LBD of Gyc76C is essential for saITP-dependent cGMP activation in cultured S2 cells, strongly suggesting that Gyc76C binds saITP directly as its ligand. Although EH acts on a different rGC, EHR (34–36), it shares with saITP an identical arrangement of four cysteine residues (C-X_2_-C-X_12_-C-X_3_-C). This structural motif may be a key determinant of ligand recognition by rGCs. In vertebrates, natriuretic peptides (NPs) and guanylins—endogenous rGC ligands—also require disulfide bonds between cysteines for receptor binding and activation (29). This suggests conserved structural features may underlie ligand recognition by rGCs across species. In a striking parallel, NPs in vertebrates also regulate water homeostasis by activating cGMP signaling and promoting natriuresis and diuresis (29).

Considering the concept of ligand/receptor co-evolution, the phylogenetic distributions of saITP and its primary receptor would be predicted to be similar, which we have confirmed for Gyc76C (Fig. 2C). Likewise, both would likely show similar loss-of-function phenotypes, and we have found this to be the case for saITP and Gyc76C, loss of either of which induces larval lethality. Despite these findings, we acknowledge that further evidence is required to conclude that Gyc76C is a *bona fide* saITP receptor. Although we did find that saITP application increased cGMP levels in S2 cells overexpressing *Gyc76C*, it did not increase cGMP levels in mammalian HEK293T cells expressing the same receptor under our assay conditions, even at same ligand concentration. Because a recent study reported that HEK293T cells expressing *Gyc76C* responded to saITP (46), it is possible that the discrepancy arises from lower sensitivity of our assay system. Nevertheless, it appears that S2 and HEK293T cells differ in their responsiveness to saITP. Moreover, saITP increased cGMP levels in *ex vivo* cultured hindguts but not in fat-body tissues, despite both tissues’ expressing *Gyc76C*. These results suggest that Gyc76C alone is insufficient for saITP signaling and that additional co-receptors or cofactors are required in a tissue-or cell-type-specific manner. Some rGCs in the nematode *Caenorhabditis elegans* function as heterodimers (47), raising the possibility that Gyc76C may likewise form functional heterodimers with other rGCs. In the vertebrate retina, guanylate cyclase-activating proteins act as essential cofactors for rGC activation (48), suggesting that similar accessory proteins may be required for Gyc76C function. It is also possible that saITP binds directly to a non-rGC receptor that then itself interacts with Gyc76C to transduce downstream cGMP signals. Indeed, some CHH-family peptides, including saITP, reportedly induce production of not only cGMP but also cAMP as second messengers (24, 41), suggesting the involvement of receptors of other families.

In conclusion, our study demonstrates that saITP functions as a neuroendocrine factor regulating systemic water balance through Gyc76C in the hindgut. Our findings uncover a brain-hindgut axis critical for osmoregulation, and they provide a conceptual framework for understanding the evolution and function of neuroendocrine water balance regulation in terrestrial arthropods.

## Materials and Methods

### Fly stocks

The following lines were obtained from the Bloomington *Drosophila* Stock Center (BDSC, Indiana University, USA): *Cg-GAL4* (stock #7011), *CG30340^attP^*(#84473), *Df(2R)ED4061* (#9068; used as a *ITP Df* strain), *Df(3L)Exel6135* (#7614; used as a *Gyc76C Df* strain), *R12G03-GAL4* (#48522), *R57C10 (nSyb)-GAL4* (#90854), *TkR99D^attP^* (#84581), *TubP-GAL4* (#5138), *TubP-GAL80^TS^* (#7019), *UAS-CG3216 RNAi* (#55895), *UAS-CG31183 RNAi* (#56937), *UAS-CG33958 RNAi* (#30507), *UAS-CG34357 RNAi* (#28524), *UAS-EHR* (*CG10738*) *RNAi #1* (#28580; unless otherwise noted, this strain was used in all *EHR* RNAi experiments), *UAS-EHR* (*CG10738*) *RNAi #2* (#57318), *UAS-Gyc76C RNAi #1* (#28660; unless otherwise noted, this strain was used in all *Gyc76C* RNAi experiments), *UAS-Gyc76C RNAi #2* (#57315), *UAS-mCD8::GFP* (on 3rd chromosome; #5130), *UAS-mCD8::GFP* (on 2nd chromosome; #5137), and *w^1118^* (the control strain; #5905).

*UAS-Dicer-2* (stock #60009) and *UAS-ITP shRNA* (#330029; unless otherwise noted, this strain was used in all *ITP* RNAi experiments) were obtained from the Vienna *Drosophila* Resource Center (VDRC, Austria). *Myo1A-GAL4* (stock #112001) and *NP1093-GAL4* (#103880) were obtained from the Kyoto Stock Center (Department of *Drosophila* Genomics and Genetic Resources, Kyoto Institute of Technology, Japan). *UAS-Gyc32E RNAi* (stock #HMJ23918) was obtained from the National Institute of Genetics (NIG, Japan). *byn-GAL4* (49) was obtained from Kenji Matsuno (Osaka University). *tsh(c724)-GAL4* (50) and *Uro-GAL4* (51) were obtained from Kenneth V. Halberg (University of Copenhagen). *Gyc76C::T2A::GAL4* was generated in-house (52).

*TM2/TM6B, Dfd-EGFP* was obtained from Takashi Nishimura (Gunma University). *Gyc76C^1^*, *ITP^SK2-4^*, *saITP^AW1^*, *saITP::2A::GAL4*, *UAS-RedcGull*, and *UAS-tethered-saITP* were generated in this study as described below.

All fly stocks were maintained on standard fly food containing 5.5 g agar (Daishin #P700), 100 g glucose (Showa Sangyo), 40 g dry yeast (Asahi Beer #HB-P02), 90 g cornmeal (Sunny Maize #Yellow-No.4M), 3 mL propionic acid (Nacalai Tesque #29018-55), 3.5 mL 10% butylparaben (in 70% ethanol) (Nacalai Tesque #06327-15), and 1 L of water. Unless otherwise noted, all experiments were conducted under non-crowded conditions at 25 °C and a 12h:12h light:dark cycle. Flies carrying the *TubP-GAL80^TS^* transgene, which encodes a temperature-sensitive inhibitor of GAL4 activity, were reared at 17 °C until the second instar and then transferred to 29 °C immediately after third instar ecdysis to induce RNAi from third instar. Unless otherwise specified, the offspring from crosses between *w^1118^* and either a mutant strain, a *GAL4* line, or a *UAS-RNAi* line were used as controls.

### Cell lines

S2 cells were maintained in 25-cm^2^ cell-culture flasks (NEST #707003) in a humidified incubator at 25 °C in Schneider’s *Drosophila* medium (SDM; Thermo Fisher Gibco #21720024) containing 10% fetal bovine serum (FBS; Thermo Fisher Gibco #10270-106) and 1% Penicillin-streptomycin solution (P/S; Fujifilm Wako #168-23191).

HEK293T cells obtained from Hiroaki Daitoku (University of Tsukuba) were maintained in Dulbecco’s Modified Eagle Medium (DMEM; Nacalai Tesque #08456-36) containing 10% FBS (Gibco #F7524) and 1% P/S (Nacalai Tesque #09367-34) in 100-mm Type-I-collagen-coated dishes (Iwaki #4020-010-MYP) in a humidified incubator at 37 °C under 5% CO_2_. Before seeding the cells on new plates, they were washed with Dulbecco’s phosphate-buffered saline (D-PBS; Nacalai Tesque #11482-15) and treated with 0.02% EDTA (Nacalai Tesque #15130-95).

### Generation of *ITP* mutants

The *ITP* mutant allele (*ITP^SK2-4^*) disrupting all splice variants was generated using the CRISPR/Cas9 as previously described (52). A 20-bp gRNA target sequence, 5’-GCGCAGCAACTTCTTCGACC-3’, was designed around the first coding exon of *ITP*. Forward and reverse 24-bp oligonucleotides with 20-bp target sequences were annealed to generate a double-stranded DNA molecule with 4-bp overhangs on each end. This was then inserted into a *BbsI*-digested pBFv-U6.2 vector (52). A transgenic strain carrying the vector was established through phiC31-mediated site-specific integration. The gRNA strain was then crossed to a *nos-Cas9* strain, and their progeny were screened by genomic PCR and sequencing for the presence of a frameshift-causing indel. We chose one such line for further analysis and named it *ITP^SK2-4^*. This line carries a frameshift mutation, a 2-bp deletion in the *ITP* exon as follows: 5’-GAAGCGCAGCAACTTCTT--ACCTGGAGTGCAAGGGCA-3’, wherein one hyphen denotes a single-nucleotide deletion.

### Generation of *saITP* isoform-specific mutants

The *saITP* isoform-specific mutant (*saITP^AW1^*) was generated using CRISPR/Cas9 as previously described (52). A 20-bp gRNA target sequence, 5’-GAACACAGGCAAGAATGCTT-3’, was designed around the initiation codon of the *saITP* coding DNA sequence (CDS; Fig. S6). Forward and reverse 24-bp oligonucleotides with 20-bp target sequences were annealed to generate a double-stranded DNA molecule with 4-bp overhangs on each end. This was then inserted into a *BbsI*-digested pBFv-U6.2 vector (52). The resulting *saITP* isoform-specific gRNA vector was injected into *w^1118^; nos-Cas9/TM6B* fly embryos (WellGenetics, New Taipei City, Taiwan). Surviving G_0_ males were crossed to 2nd chromosome *Sp/SM6a* virgin females, and their progeny were screened by genomic PCR and sequencing for the presence of frameshift-inducing indels. We chose one line for further analysis and named it *saITP^AW1^*. This is a frameshift mutant carrying a 14-bp deletion starting at the second nucleotide of the *saITP* CDS, which disrupts the entire saITP amino-acid sequence (Fig. S6).

### Generation of *saITP::T2A::GAL4* flies

The *saITP::T2A::GAL4* line was generated by CRISPR-mediated targeted integration of *T2A::GAL4* immediately in front of the stop codon of *saITP* as previously described (53). A donor vector for targeted integration was generated by assembling the left and right homology arms and the *T2A::GAL4/3xP3-RFP* cassette in a single enzymatic reaction using In-Fusion (Takara). The left homology arm was amplified by the following primer set:

F: 5’-GCTTGATATCGAATTCACGCAAGATCGCAATTTGTGCAAGACAC-3’ R: 5’-AGTTGGGGGCGTAGGCTTGCGACCCAGGGTATCGCGCCATTCG-3’.

The right homology arm was amplified by the following primer set:

F: 5’-TAGTATAGGAACTTCAGTGCGATTCTCTGGGATTTTCTGGCAC-3’

R: 5’-CGGGCTGCAGGAATTCCCGCAATCAAATTCGCTGTCGATTCCTC-3’.

Underlined sequences above match the genomic DNA sequence, and the rest of the sequences are adapters for vector construction. The donor vector and a gRNA vector expressing the sequence 5’-GCAAGTAAAGTGCGATTCTC-3’ were co-injected into fertilized eggs of the *nos-Cas9* strain to induce a site-specific double-strand break and subsequent integration of the donor vector. The injected embryos were raised to adulthood, and their progeny were screened for transformants marked by eye-specific RFP fluorescence. A single transformant was crossed to a *TM6B* balancer line to establish a stock. The “floxed” *3xP3-RFP* marker was removed before use by crossing to a *TM6B, Cre-w* balancer line.

### Total RNA extraction and quantitative reverse transcription (qRT)-PCR

Whole bodies or dissected tissues were collected in 1.5 mL tubes and immediately flash-frozen in liquid nitrogen. Total RNA from bodies or tissues was extracted as previously described (55). cDNA was generated from purified total RNA using ReverTra Ace qPCR RT Master Mix with gDNA Remover (TOYOBO #FSQ-301) according to the manufacturer’s instructions. qRT-PCR was performed on a CFX Duet Real-Time PCR machine (Bio-Rad) using Taq Pro Universal SYBR qPCR Master Mix (Vazyme #Q712-02). For absolute mRNA quantification, serial dilutions of pGEM-T (Promega #A362A) plasmids containing the coding sequences of a target gene or *rp49* were used for standards. After the molar amounts were calculated, the transcript levels of each target mRNA were normalized to the levels of *rp49* in the same samples. Three separate samples were collected for each experiment, and duplicate measurements were conducted. The primers to detect *rp49* levels were previously reported (54), and the others are as follows:

*saITP*-F: 5’-ACACCGATTATGCAAGCAAGAATGC-3’ *saITP*-R: 5’-TATCGCGCCATTCGTTATACTTGTC-3’ *ITPL1*-F: 5’-ATCGACGATGAAGAGATATCGCAGC-3’ *ITPL1*-R: 5’-AGTGGTCGAAGTTAGGAGTCTAGTG-3’ *ITPL2*-F: 5’-ACAGCTGAGAATCGGTGCGAATATC-3’ *ITPL2*-R: 5’-TTAGATTTCTGTGTTCTGGCTCTGC-3’ *Gyc76C*-F: 5’-TTCTGTACGAGGATGTGTGGAGTCC-3’ *Gyc76C*-R: 5’-AGCATCTGCTCACAGCACTTGACTC-3’

### Analysis of larval survival and mortality

Larval 24-hr survival rate was calculated based on the proportion of newly hatched larvae surviving the first 24 hrs after hatching. Newly hatched larvae were transferred to small food vials at a density of 25 larvae per vial. The vials were then maintained in a humidified incubator, and the number of surviving larvae was recorded after 24 hrs. Larval mortality was calculated based on the proportion of larvae that developed to the pupal stage as previously described (55). Newly hatched larvae were transferred to small food vials at a density of 25 larvae per vial and then maintained in a humidified incubator. The number of pupae that formed in each vial was recorded.

### Immunostaining

Tissues were dissected in 1x PBS (Nacalai Tesque #27575-31), fixed with 4% paraformaldehyde (PFA; Nacalai Tesque #02890-45) in PBS (4% PFA/PBS) containing 0.3% Triton X-100 (Nacalai Tesque #35501-15) for 20 min at room temperature (RT) and then washed multiple times with PBS containing 0.3% Triton X-100 (PBST). Tissues were blocked with 2% bovine serum albumin (BSA; Sigma-Aldrich #A9647) in PBST for at least 1 hr at RT, incubated overnight at 4 °C with primary antibodies mixed in PBST containing 2% BSA, washed multiple times with PBST, incubated for 2 hrs at RT with secondary antibodies in PBST containing 2% BSA, and again washed multiple times with PBST. When appropriate, DAPI (Sigma-Aldrich #D9542) nuclear stain was diluted 1:2,000 into the secondary-antibody mixture. After washing, the tissues were mounted in Vectashield H-1000 (Vector Laboratories) and observed using a Zeiss Axio Imager M2 equipped with ApoTome.2 or a Zeiss LSM 900 confocal microscope. The specificity of the signals was established by comparison with appropriate controls.

Chicken anti-GFP (1:2,000 dilution, Abcam #ab13970) and guinea pig anti-saITP (1:1000 dilution; ref. 57) were used as primary antibodies. Alexa Fluor 488 goat anti-chicken (1:2,000 dilution, Thermo Fisher Scientific #A32931) and Alexa Fluor 488 goat anti-guinea pig (1:2,000 dilution, Thermo Fisher Scientific #A11073) were used as secondary antibodies.

### Phylogenetic tree analysis

We generated neighbor-joining trees using ClustalW (https://www.genome.jp/tools-bin/clustalw). We aligned the amino acid sequences of seven *D. melanogaster* rGCs to 58 rGCs from vertebrates (Human, *Homo sapiens*; Mouse, *Mus musculus*; Zebrafish, *Danio rerio*), holometabolous insects (yellow fever mosquito, *Aedes aegypti*; silk moth, *Bombyx mori*; red flour beetle, *Tribolium castaneum*; western honey bee, *Apis mellifera*), a hemimetabolous insect (human body louse, *Pediculus humanus*), and a crustacean (water flea, *Daphnia pulex*). Protein names and GenBank accession numbers are listed in Table S1.

### Construction of plasmids for transfection

*pUAST-Gyc76C*, *pcDNA-Gyc76C*, and *pUAST-EHR* plasmids were constructed from cDNA clones LD12174 (for *Gyc76C*) and IP14815 (for *EHR*), which were obtained from the *Drosophila* Genomic Resource Center (DGRC, Indiana University, USA). These cDNA clones were used as templates for PCR amplification using KOD Plus Neo (TOYOBO #KOD-401). The resulting PCR products were cloned into *Eco*RI- and *Xba*I-digested *pUAST* vector or *pcDNA3.0* vector using the In-Fusion HD Cloning Kit (Takara #102518) according to the manufacturer’s instructions.

*pUAST-Gyc76C-ΔLBD* was constructed from a cDNA clone generated by Twist Bioscience (San Francisco, USA). The domain organization of Gyc76C was predicted using InterPro (https://www.ebi.ac.uk/interpro/) and used to design a *Gyc76C-ΔLBD* sequence lacking the predicted LBD (glycine 48 to proline 424, numbered from the initiator methionine). The *Gyc76C-ΔLBD* sequence was cloned into *Eco*RI- and *Xba*I-digested *pUAST* vector using the In-Fusion HD Cloning Kit according to the manufacturer’s instructions. The primers used were as follows:

*pUAST-EHR*-F: 5’-AGGGAATTGGGAATTAGCAGCATTTCACGCGATTC-3’ *pUAST-EHR*-R: 5’-ACAAAGATCCTCTAGAGGACACTGCCTTAAACCTC-3’ *pUAST-Gyc76C*-F: 5’-AGGGAATTGGGAATTTCGCTGTGAGAATCCGTTGG-3’ *pUAST-Gyc76C*-R: 5’-ACAAAGATCCTCTAGCTACACTGCCTTCTCCTTAT-3’ *pUAST-Gyc76C-ΔLBD*-F: 5’-AGGGAATTGGGAATTATGACGCGTTGGCCCTTTAA-3’ *pUAST-Gyc76C-ΔLBD*-R: 5’-ACAAAGATCCTCTAGCTATACTATACTCTCACGAT-3’ *pcDNA3-Gyc76C*-F: 5’-AGTGTGCTGGAATTTCGCTGTGAGAATCCGTTGG-3’ *pcDNA3-Gyc76C*-R: 5’-ATAGGGCCCTCTAGCTACACTGCCTTCTCCTTAT-3’

### Generation of *ITP* double-stranded (ds) RNA

For dsRNA preparation, the 20-bp T7 promoter sequence (underlined) was attached to the 5′ end of the *ITP*-specific PCR primers as follows:

F: 5’-<u>GGATCCTAATACGACTCACTATAGG</u>ACCACAACCAGCGATAAGCATCCG-3’ R: 5’-<u>GGATCCTAATACGACTCACTATAGG</u>AACAACTGGTAGCAGTCCTCGCAG-3’

Whole-body cDNA was used as a PCR template. The resulting PCR product carrying the T7 promoter sequences was used as a template for *in vitro* transcription using the RiboMAX Express RNAi System (Promega #P1700) according to the manufacturer’s instructions.

### Generation of synthetic peptides

Mature *Drosophila* saITP peptide, comprising 73 amino acid residues (NH_2_-SNFFDLECKGIFNKTMFFRLDRICEDCYQLFRETSIHRLCKQECFGSPFFNACIEALQLHEEMDKYNEWRDTL-CONH_2_), was synthesized using fluorenylmethyloxycarbonyl (Fmoc) chemistry and microwave-assisted solid-phase peptide synthesis using a Initiator+ Alstra automated peptide synthesizer (Biotage, Uppsala, Sweden) as previously described (57). The refolding of saITP was performed as previously described (58). High-performance liquid chromatography (HPLC) was used to confirm a purity >95% for the synthesized saITP. Lyophilized saITP was weighed on an analytical balance (AP125WD; Shimadzu, Kyoto, Japan). Mature *Drosophila* NPLP1-VQQ peptide (NH_2_-NLGALKSSPVHGVQQ-COOH) was synthesized by Eurofins Genomics (Tokyo, Japan).

### Transfection and cGMP assay in culture cells

Just prior to transfection, S2 cells were seeded at a density of 5×10^5^ cells/mL into 35-mm cell-culture dishes (Greiner #627160) in 2 mL of SDM containing 10% FBS and 1% P/S. S2-cell transfection was performed using the Effectene transfection reagent (QIAGEN #301425) according to the manufacturer’s instructions. Each dish was transfected with 0.3 μg *pUAST* empty vector (as a control) or *pUAST-rGC* (*pUAST-Gyc76C*, *pUAST-EHR*, or *pUAST-Gyc76C-ΔLBD*) along with 0.1 μg of *pActin-GAL4*. For the *ITP* RNAi experiments, 10 μg of *ITP dsRNA* was added to each dish at the time of transfection and again every 24 hrs afterward. The cells were incubated for three days after transfection. Then, 500 μL/well of transfected cells were transferred to a 24-well plate (TPP #92424). After 4 hrs of incubation, 50 μL of the medium in each well was removed and replaced with 50 μL of 10-mM 3-isobutyl-1-methylxanthine (IBMX) (Adipogen Life Sciences #AG-CR1-3512-M500) to reach the final concentration of 1 mM. After a 30-min incubation, 50 μL of the medium in each well was removed and replaced with 50 μL of SDM (as a vehicle control) or synthetic mature saITP stock, for the final concentration of 100 nM, or mature Nplp1-VQQ (at the final concentration of 1 μM) in SDM. After another 30-min incubation, the medium was removed and replaced with 500 μL of 0.1-M HCl to lyse the cells. After a 10-min incubation at RT, the cell lysates were collected and stored at -80 °C.

Just prior to transfection, HEK293T cells were seeded at a density of 5×10^5^ cells/mL into 35-mm collagen-coated dishes (Iwaki #4000-010-MYP) in 2 mL of DMEM containing 10% FBS and 1% P/S. HEK293T cells were transfected with 0.4 μg/dish of *pcDNA3.0* empty vector (as control) or *pcDNA-Gyc76C*, using the Effectene transfection reagent according to the manufacturer’s instructions. Three days after transfection, the transfected cells were washed with D-PBS, treated with 0.02% EDTA, and resuspended in 2 mL of DMEM containing 10% FBS and 1% P/S. Cells were re-seeded at 500 μL /well in a 24-well collagen-coated plate (Iwaki #4820-010). After a 4-hr incubation, 50 μL of the medium in each well was replaced with 50 μL of 10 mM IBMX to attain the final concentration of 1 mM. After a 30-min incubation, another 50 μL of the medium in each well was replaced with 50 μL of DMEM (as a vehicle control) or mature saITP peptide (at the final concentration of 100 nM) in DMEM. Finally, after a 30-min incubation, the medium was removed, and 500 μL of 0.1 M HCl was added to each well to lyse the cells. After a 10-min incubation at RT, the cell lysates were collected and stored at -80 °C.

After cell lysis, cGMP levels were determined using a cGMP ELISA Kit (Cayman #581021) according to the manufacturer’s instructions, with indicator levels measured using a Multiskan GO spectrophotometer (ThermoFisher Scientific).

### Generation of *Gyc76C* mutant flies

The *Gyc76C^1^* mutant allele was generated using the CRISPR/Cas9 as previously described (52). Pairs of gRNA target sequences (20 bp) were designed near the *Gyc76C* translational start (T1: 5’-GGATTGTGTTGGCGTGGACA-3’) and stop sites (T2: 5’-GAAGAAGGAGCAGGATGACC-3’). Forward and reverse 24-bp oligonucleotides with 20-bp target sequences were annealed to generate a double-stranded DNA with 4-bp overhangs on each end. This was then inserted into *BbsI*-digested pBFv-U6.2 or pBFv-U6.2B vector (52). To construct a double-gRNA vector, the first gRNA (T1) was cloned into pBFv-U6.2 (named pBFv-U6.2-T1). Then, the second gRNA (T2) was cloned into pBFv-U6.2B (named pBFv-U6.2B-T2). A fragment containing the U6 promoter and the first gRNA was cut from pBFv-U6.2-T1 and ligated into pBFv-U6.2B-T2. This *Gyc76C* double-gRNA vector was then injected into *yw; nos-Cas9[attP40]/CyO* fly embryos (BestGene Inc, CA, USA). Surviving G_0_ males were divided into 10 groups and crossed *en masse* to 3rd chromosome balancer virgins *(TM2/TM6B, Dfd-EGFP)*. Five individual males were isolated from the progeny of each of these ten crosses and crossed independently to *TM2/TM6B, Dfd-EGFP* virgins to establish independent isogenized lines. Among the resulting 50 lines, those with homozygous lethality were isolated. To confirm deletions at the *Gyc76C* locus, we randomly selected 5 lines and performed genomic PCR and sequencing. We confirmed complete deletions in all five lines and chose one line for further analysis, naming it *Gyc76C^1^*. *Gyc76C^1^* has a 7,403-bp deletion that includes the entire *Gyc76C* coding DNA sequence (CDS).

### Generation of *UAS-RedcGull* transgenic flies

A codon-optimized *RedcGull* cassette (Fig. S7) comprising an *Eco*RI cut site, *Drosophila* Kozak sequence, *Drosophila*-optimized RedcGull coding sequence, and *Xba*I cut site was synthesized and sequence-verified by GeneArt Services (Thermo Fisher Scientific, MA, USA). This cassette was cloned into *Eco*RI-and *Xba*I-digested *pUAST-attB* vector. This *pUAST-attB-RedcGull* construct was inserted into the VK00018 *attP* site on chromosome 2R via phiC31 integrase-mediated transgenesis (BestGene Inc, CA, USA).

### cGMP imaging analysis of *ex vivo* cultured tissues

cGMP imaging analysis of *ex vivo* cultured tissues was performed using flies expressing *UAS-RedcGull* under the control of tissue-specific *GAL4* drivers (*i.e.*, *byn-GAL4* for the hindgut and *Cg-GAL4* for the fat body). Tissues were dissected from wandering third instar in SDM with 10% FBS and 1% P/S and then washed with SDM. The dissected tissues were kept in a glass-bottomed dish (35×10 mm, IWAKI, #3910-035) with insect pins (ϕ0.10 mm, Ento Sphinx Insect Pins) and silicone grease (Beckman) covered with 20 μL of SDM. Live imaging was performed at RT using a Zeiss LSM 900 confocal microscope. RedcGull was excited with a 555-nm laser. Time-lapse images were acquired every 20 seconds for 420 seconds. Then, 120 seconds after starting live-imaging, 80 μL of SDM with or without 100 nM synthetic saITP was added to the tissue. Mean fluorescence intensities were measured along the time axis using Fiji (ImageJ2, v2.16.0) and R (v4.4.2).

### Generation of *UAS-tethered-saITP* transgenic flies

Because saITP is amidated on its C-terminal end, we chose to create a membrane-tethered saITP by fusing a transmembrane domain to the N terminus of the predicted secreted form of saITP. To do this, we adapted a technique first described for constructing tethered-Bursicon constructs (59, 60). A transmembrane domain known to function in a “type-2” orientation (*i.e.*, with an intracellular N terminus and extracellular C terminus) was fused between an N-terminal mTagBFP2 fluorophore with an extracellular spacer sequence and the saITP domain (Fig. S8). Since saITP relies on disulfide bridges to retain proper structure, no other cysteine residues were included in the extracellular sequence to promote proper folding. The amino-acid sequence (Fig. S8) of the construct was reverse-translated and codon-optimized for *D. melanogaster*.

A codon-optimized *tethered-saITP* cassette (Fig. S8) comprising an *Eco*RI cut site, *Drosophila* Kozak sequence, *Drosophila*-optimized *tethered-saITP* coding sequence, and *Xba*I cut site was synthesized and sequence-verified by GeneArt Services (Thermo Fisher Scientific, MA, USA). This cassette was cloned into *Eco*RI-and *Xba*I-digested *pUAST-attB* vector. Then, the *pUAST-attB tethered-saITP* plasmid was inserted into the *attP40* site on chromosome 2L via phiC31 integrase-mediated transgenesis (BestGene Inc, CA, USA).

### *Ex vivo* fluid-reabsorption assay

Water reabsorption in the hindgut was assessed using an *ex vivo* culture protocol modified slightly from one previously described (44, 45, 61). Embryos were collected on apple-juice plates with a small amount of yeast paste for 4 hrs at 25 °C. The plates were then transferred to 18 °C and incubated for an additional 48 hrs, after which newly hatched first-instar larvae were collected and transferred to standard fly-food vials (30-35 per vial). Around 100 hrs after larval selection, all the vials were shifted to 29 °C. Large actively feeding mid-third instar larvae were used for reabsorption assays. Larvae were thoroughly washed twice with MilliQ water. Then, in SDM, the cuticle and fat body were carefully removed without disturbing the CNS complex, gut, Malpighian tubules, or hindgut. The tracheal trunks were cut to facilitate mounting. It is important to note that the posterior cuticle was carefully removed apart from the posterior spiracles, while the anterior cuticle around the mouthparts was retained to enable pinning. After removing residual SDM, the dissected organ complex was gently transferred to a paraffin-wax dish filled with water-saturated paraffin oil (Sigma-Aldrich #18512). To preserve the foregut, midgut, hindgut, and tubules, the complex was gently stretched and pinned at the anterior cuticle and posterior spiracles. Any excess SDM was removed using a pipette. Two drops of assay medium [1:1 premixed HL3.1 saline (62) and SDM supplemented with 100 µM of the red dye amaranth (Sigma-Aldrich #A1016) in DMSO] were added under paraffin oil. One drop (2 µL) was placed around the midgut and tubules, and a second smaller 0.3 µL drop—either with or without 100 nM saITP—was placed near the hindgut (while avoiding the posterior spiracles). The tubules were gently drawn away from the large drop and wrapped around a pin using fine forceps. The circumference of the smaller droplet was measured using an eyepiece graticule (Leica, Germany). Specimens were excluded from further analysis if they lacked any of the following: (1) gut peristaltic contractions within one min of adding the assay medium, (2) pigmentation of the tubules (indicating a lack of amaranth uptake), or (3) enlargement of the posterior droplet. These were rare under control conditions (*byn^ts^*>*w^1118^* with no saITP application). One hr after droplet placement, changes in the circumference of the initial and final drops were converted to volumes using the following equation: V = (π x d^3^)/6. Then, the rate of absorption (nl/min) was calculated as Δv/Δt, where Δv is the change in volume (nl) and Δt is the duration (min) of the experiment.

### Quantification of body water volume

Larval wet weight was measured by individually weighing actively feeding third instar larvae in 0.2 mL PCR tubes on an ultra-microbalance (Sartorius, SE2). After weighing, the larvae were instantly killed on dry ice, transferred to a 60 °C oven, and dried for 36 hrs. The dried larvae were re-weighed to assess the amount of water lost. Body water volume was calculated as percent water using the following formula: 100 x ([wet weight] - [dry weight]) / [wet weight].

### Hemolymph collection and osmolality measurement

For hemolymph collection, third instar larvae were rinsed in water, gently blotted dry, and placed in a clean glass dish on ice. The cuticle was nicked with fine tweezers, carefully avoiding damaging internal tissues such as the gut to prevent contamination. The exuding hemolymph was immediately collected by pipette into pre-chilled Eppendorf tubes on ice, and hemolymph from multiple larvae was pooled. To limit melanization, samples were heat-treated at 70 °C for 5 min before being frozen. To purify the hemolymph for osmolarity measurements, samples were thawed and centrifuged to remove circulating cells (top speed, 5 min) at 4 °C, and the clarified supernatant was used for osmometry. Osmolarity was measured by using 10 µL of hemolymph per sample on a vapor-pressure osmometer (VAPRO model 5600, ELITechGroup) calibrated to manufacturer standards. Sampling and loading were performed rapidly to minimize evaporation; each biological replicate corresponded to one pooled hemolymph sample.

### Quantification of body weight

After puparium formation, intact pupae were carefully cleaned using distilled water and a paintbrush and then completely dried on paper towels. Clean, surface-dry (not dehydrated) pupae were individually weighed on an ultra-microbalance (Sartorius, SE2).

## Acknowledgements

We thank the Vienna *Drosophila* Resource Center, the Bloomington *Drosophila* Stock Center, the National Institute of Genetics Fly Stock Center, the Kyoto Stock Center, the Transgenic RNAi Project at Harvard Medical School, K. Matsuno, T. Nishimura, K.V. Halberg, and H. Tanimoto for fly stocks; H. Daitoku for cell lines; FlyBase for providing comprehensive and invaluable resources on *Drosophila*; K.V. Halberg for access to the vapor-pressure osmometer; K.V. Halberg, M. Gáliková, and D. Žitňan for helpful discussions; and Write Labs (https://write-labs.com) for help with editing the manuscript. This study was supported by the Japan Science and Technology Agency (JST) FOREST Program (JPMJFR224M), the Japan Society for the Promotion of Science (JSPS) KAKENHI Grant-in-Aid for Scientific Research (B) (25K02307) to N.O., the JSPS KAKENHI Grant-in-Aid for JSPS Fellow (24KJ0466), the JST SPRING Program (JPMJSP2124) to A.W., and the Independent Research Fund Denmark (DFF) (4283-00022B) to K.R. A.W. is the recipient of a fellowship from the Japan Society for the Promotion of Science.

## Author Contributions

N.O. conceived the study. A.W., T.K., O.K., M.J.T., K.R., R.N., and N.O designed the experiments and interpreted the data. A.W. performed most of the experiments and analyzed the data with contributions from T.K., O.K., M.J.T., K.R., R.N., and N.O. S.K. contributed to the generation of *ITP^SK2-4^* and *saITP::T2A::GAL4*. A.W. and N.O. contributed to the generation of *saITP^AW1^*. R.M., N.O., and N.Y. contributed to the generation of *Gyc76C^1^*. T.K., O.K., M.J.T., and K.R. contributed to the generation of *UAS-RedcGull* and *UAS-tethered-ITP*. M.F., E.I-U., and K.U. contributed to the synthesis of the saITP peptide. T.Y. contributed to the generation of the anti-saITP antibody. A.W. and N.O. wrote the manuscript with contributions from T.K., O.K., M.J.T., H.T., S.K., N.Y., K.U., K.R., and R.N. N.O. supervised the project.

## Corresponding authors

Correspondence to Naoki Okamoto.

