## Supplemental Figures S1 to S8, and Table S1 for "Ion transport peptide regulates body water balance via a receptor guanylyl cyclase in the *Drosophila* hindgut"

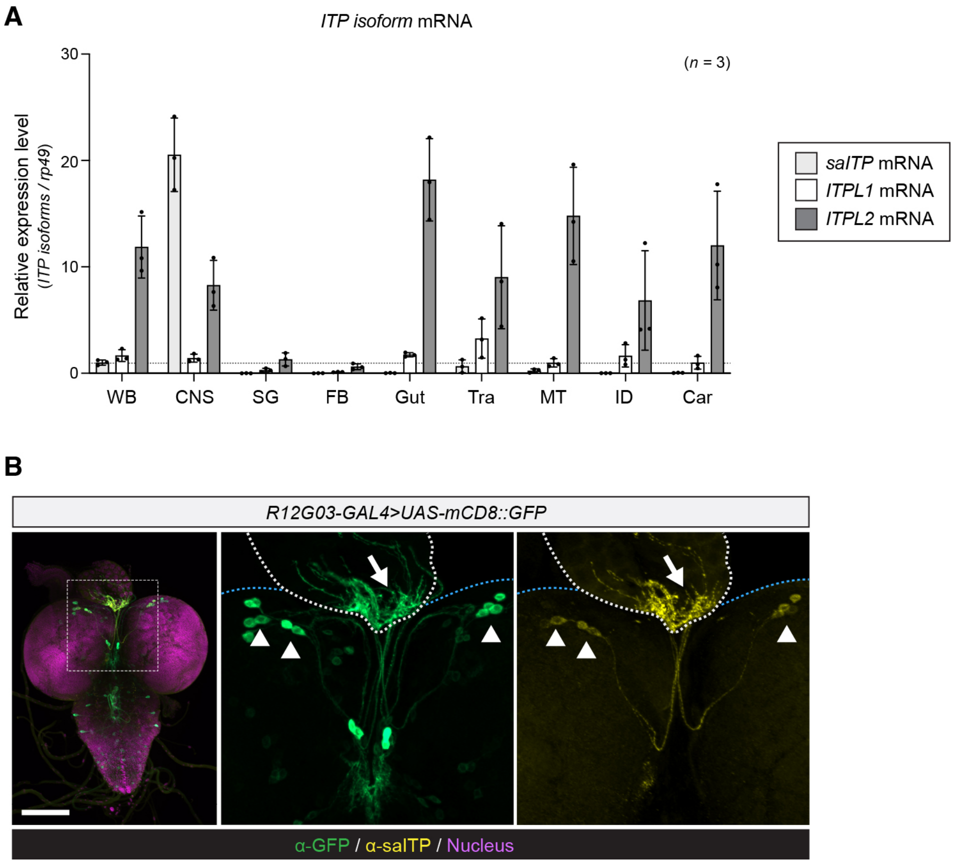


**Figure S1: Expression pattern of the *ITP* isoforms and *R12G03-GAL4*.**

**A**, Relative expression of the *ITP* isoforms *saITP*, *ITPL1*, and *ITPL2* in various tissues, as assessed by RT-qPCR. WB, whole body; CNS, central nervous system; SG, salivary glands; FB, fat body; Tra, tracheae; MT, Malpighian tubules; ID, imaginal discs; Car, carcass. For absolute quantification, serial dilutions of plasmids carrying the respective cDNAs were used for standards. After the molar amounts were calculated, the transcript levels for each *ITP* isoform were normalized to *rp49*. Values are shown relative to WB *saITP* expression. **B**, *R12G03-GAL4* expression pattern (green) and saITP immunoreactivity (yellow) in the CNS dissected from wandering third instar larvae. Nuclei were stained with DAPI (magenta). *R12G03-GAL4* was crossed to *UAS-mCD8::GFP*. The right image shows a magnified view of the region within the white-dotted square. The arrowheads indicate the *ipc-1* neurons, and the arrow indicates the CC. Scale bars, 100 μm. Data are presented as means ± SD. Sample sizes (*n*) are shown in each graph.

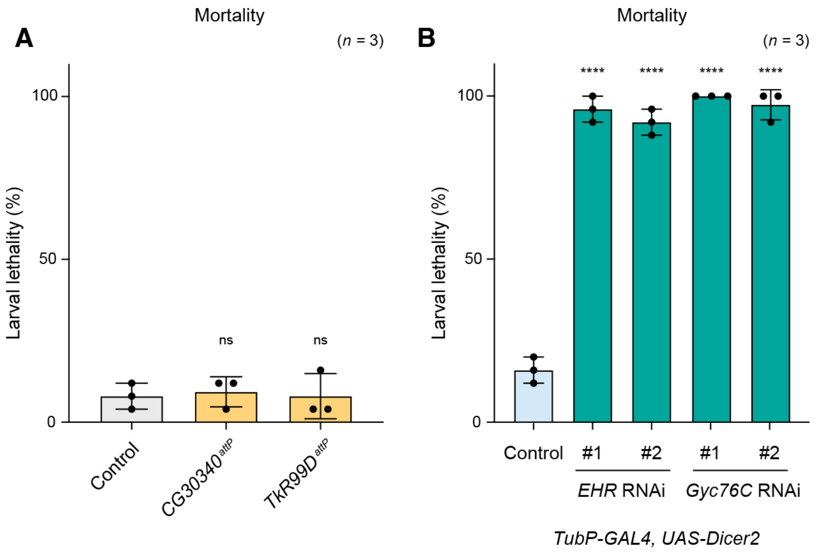


**Figure S2: Larval lethality in mutants and candidate saITP receptor knockdown animals.** **A**, Larval mortality for the control (*w*$^{1118}$), *CG30340* homozygous mutants (*CG30340*$^{attP}$), and *TkR99D* homozygous mutants (*TkR99D*$^{attP}$). **B**, Larval mortality upon ubiquitous knockdown of *EHR* or *Gyc76C*. *TubP-GAL4>UAS-Dicer-2* was used as a ubiquitous driver. Two independent RNAi lines were used to confirm the phenotype. Larvae were reared at 25 individuals per vial, and the data represent the results of three independent experiments. Data are presented as means ± SD. Sample sizes ($n$) are shown in each graph. ****$p < 0.0001$, as assessed by one-way ANOVA with Dunnett's test for multiple comparisons to control. ns: not significant ($p > 0.05$).

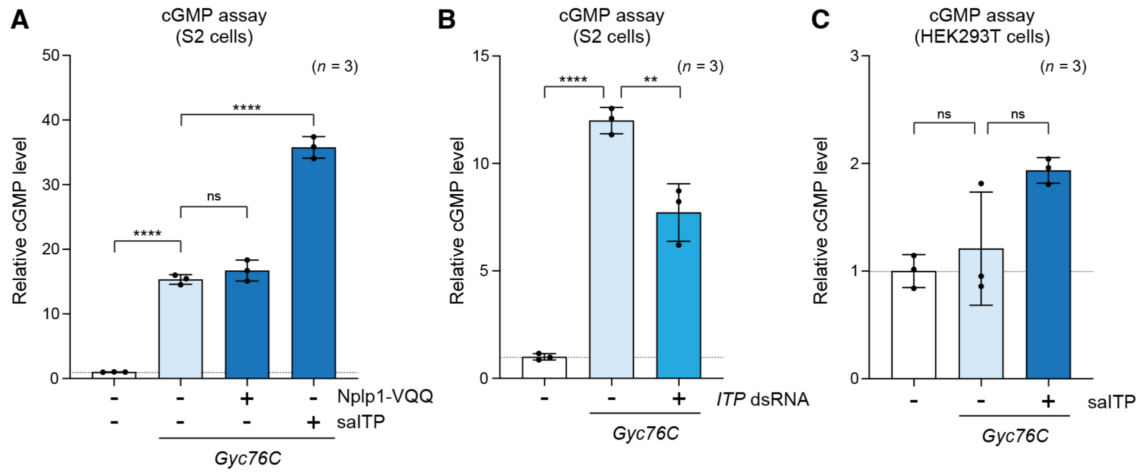


**Figure S3: *In vitro* cGMP assay using S2 cells and HEK293T cells.**

**A**, Intracellular cGMP levels in S2 cells expressing *Gyc76C* with (+) or without (−) the addition of synthetic Nplp1-VQQ at the final concentration of 1 µM. Synthetic saITP at the final concentration of 100 nM was added as a positive control. **B**, Effect of *ITP* knockdown on intracellular cGMP levels of S2 cells expressing *Gyc76C*. *pActin-GAL4* was used to express *Gyc76C* in S2 cells (**A** and **B**). **C**, Intracellular cGMP levels in HEK293T cells expressing *Gyc76C* with (+) or without (−) the addition of synthetic saITP. Values are relative to the control (−). Data are presented as means ± SD. Sample sizes ($n$) are shown in each graph. $^{**}p < 0.01$, $^{****}p < 0.0001$, as assessed by one-way ANOVA with Tukey's test for multiple comparisons. ns: not significant ($p > 0.05$).

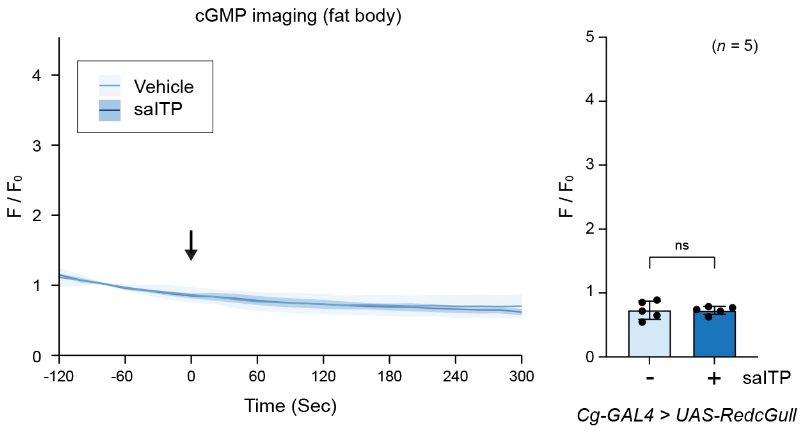


**Figure S4: *Ex vivo* cGMP assay using the fat body.**

RedcGull fluorescence intensity in the fat body of wandering third instar larvae with (+) or without (−) the addition of synthetic saITP. The left graphs show temporal changes in RedcGull fluorescence intensity, while the right graphs show a quantification of RedcGull fluorescence intensity at 120 seconds after the stimulation. The average fluorescence intensity immediately before the addition of the reagent was set to $F_0$, and the values at all time points were normalized to $F_0$. *Cg-GAL4* was used as a fat-body-specific driver.

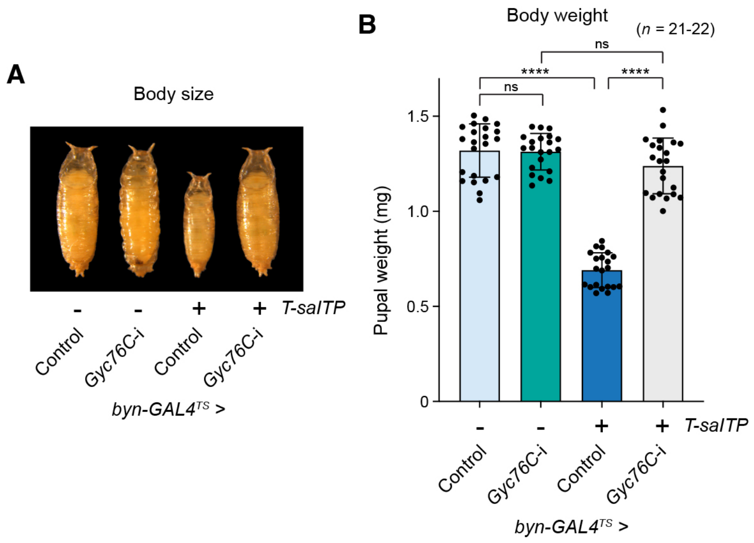


**Figure S5: Hindgut-specific expression of *T-saITP* severely impaired body growth.**

**A** and **B**, Representative images (**A**) and body weight (**B**) of control and *Gyc76C* RNAi (*Gyc76C-i*) pupae with (+) or without (−) expression of *tethered-saITP* (*T-saITP*). In all *Gyc76C*-knockdown experiments, larvae were reared at 18 °C during the early larval stages and shifted to 29 °C from 0 hr after third instar ecdysis to induce RNAi from the third instar. *byn-GAL4*$^{TS}$ was used as a hindgut-specific driver. Data are presented as means ± SD. Sample sizes (*n*) are shown in each graph. ****$p < 0.0001$, as assessed by one-way ANOVA with Tukey's test for multiple comparisons. ns: not significant ($p > 0.05$).

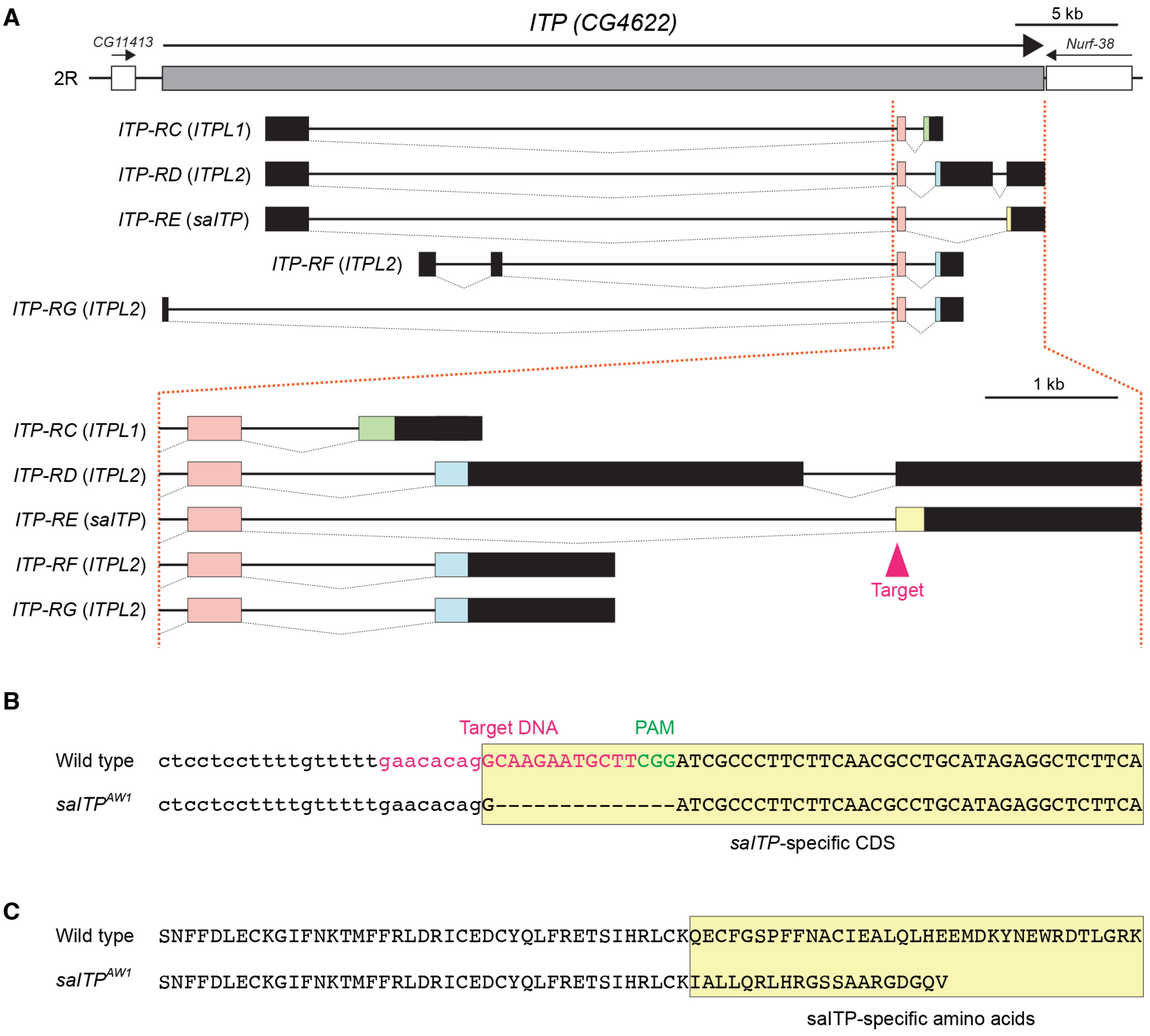


**Figure S6: Generation of *saITP*-specific mutants.**

**A**, Schematic representation of the *ITP* locus and guide RNA (gRNA) target. The protein-coding DNA sequence (CDS) for all *ITP* isoforms and untranslated regions are represented by colored open boxes and filled boxes, respectively. Neighboring genes are represented by white boxes. Arrows indicate the orientation of the *ITP* gene. The gRNA target sequences (20 bp) were designed around the start point of the *saITP* CDS. **B**, DNA sequence for $saITP^{AW1}$. The wild-type sequence appears above. The gRNA target sequence appears in red, and the neighboring NGG protospacer adjacent motif (PAM) sequence is shown in green. Deleted residues are shown as dashes. $saITP^{AW1}$ has a 14-bp deletion beginning with the second nucleotide of the *saITP* CDS. **C**, Amino-acid sequence of $saITP^{AW1}$. The entire saITP amino-acid sequence is disrupted in $saITP^{AW1}$ because the 14-bp deletion created a frameshift.

*Eco*RI Kozak Start

```
GAATTCGCCACCATGCTGGTGTCCAAGGGCGAAGAGGATAACATGGCCATCATCAAAGAGTTCATGCGCTTCAAGGTGCACATGG
AAGGCAGCGTGAACGGCCACGAGTTCGAGATTGAAGGCGAAGGCGAGGGACGCCCATACGAGGCCTTTCAAACCGCCAAGCTGAA
AGTGACCAAAGGCGGCCCACTGCCGTTCGCCTGGGATATTTTGAGCCCGCAGTTTATGTACGGCAGCAAGGCCTACATCAAGCAC
CCCGCCGATATTCCCGATTACTTCAAGCTGAGCTTCCCCGAGGGATTCCGCTGGGAGCGCGTGATGAATTTCGAGGATGGCGGCA
TCATCCACGTGAACCAGGATAGCAGTCTGCAGGATGGCGTGTTCATCTACAAAGTGAAGCTGCGCGGCACGAACTTCCCACCAGA
TGGCCCAGTGATGCAGAAAAAGACCATGGGCTGGGAAGCCCTGAGCTTCAGCCTGGATCTGGGCCTGCACATCCACGGACTGATT
AGCGCCGATCGCTACAGCCTGTTCCTCGTGTGCGAGGATAGCTCCAAGGATAAGTTCCTGATCAGCCGCCTGTTCGATGTGGCCG
AGGGAAGTACACTGGAAGAGGCCAGCAACAACTGCATCCGCCTGGAATGGAACAAGGGCATCGTGGGACATGTGGCCGCCTTCGG
AGAGCCCCTGAATATCAAGGATGCCTACGAGGATCCCCGCTTCAACGCCGAGGTGGACCAGATTACCGGCTACAAGACCCAGAGC
ATCCTGTGCATGCCCATCAAGAACCACCGCGAAGAGGTTGTGGGAGTCGCCCAGGCCATCAACAAGAAGAGTGGAAACGGCGGCA
CCTTCACCGAAAAGGATGAGAAGGATTTCGCCGCCTACCTGGATCGCCTGAAGAAAGAACTGCAAGAGTTCGTGTCCGAGCGCAT
GTACCCAGAGGATGGTGCCCTGAAGTCCGAGATCAAGAAGGGCCTGCGCCTGAAGGATGGTGGACATTACGCCGCCGAAGTCAAG
ACCACCTACAAGGCCAAGAAACCCGTGCAACTGCCAGGCGCCTACATCGTGGATATCAAGCTGGACATCGTGTCCCACAACGAGG
ATTACACCATCGTGGAACAGTGCGAGCGCGCCGAAGGACGTCATAGTACAGGCGGAATGGATGAGCTGTACAAGTAGTCTAGA
```

Stop *Xba*I

**Figure S7: Construction of a codon-optimized *RedcGull* cassette.**

To enable real-time monitoring of cGMP changes in *Drosophila* cells and tissues, the coding sequence of the RedcGull sensor (43) was adapted for use under GAL4 control. The resulting cassette comprises an *Eco*RI site, a *Drosophila* Kozak sequence, a *Drosophila*-optimized *RedcGull* coding sequence, and an *Xba*I site.

**A**

mTagBFP2 - Transmembrane Domain - Linker - saITP (amidation sequence)

```
MVSKGEELIKENMHMKLYMEGTVDNHHFKCTSEGEGKPYEGTQTMRIKVVEGGPLPFAFDILATSFLYGSKTFINHTQGIPDFFK
QSFPEGFTWERVTTYEDGGVLTATQDTSLQDGCLIYNVKIRGVNFTSNGPVMQKKTLGWEAFTETLYPADGGLEGRNDMALKLVG
GSHLIANAKTTYRSKKPAKNLKMPGVYYVDYRLERIKEANNETYVEQHEVAVARYCDLPSKLGHKLNGGSMSTESMIRDVELAEE
ALPKKTGGPQGSRRCLFLSLFSFLIVAGATTLFCLLHFGVIGSGNGNGNGNGNGNGNGNGNGNEQKLISEEDLGNGNGNGSLESN
FFDLECKGIFNKTMFFRLDRICEDCYQLFRETSIHRLCKQECFGSPFFNACIEALQLHEEMDKYNEWRDTLGRK*
```

**B**

*Eco*RI Kozak Start

```
GAATTCGCCACCATGGTTTCCAAGGGCGAAGAACTGATCAAAGAAAACATGCACATGAAGCTGTACATGGAAGGCACCGTGGATA
ACCACCACTTCAAGTGCACCAGCGAAGGCGAGGGCAAGCCATATGAGGGAACCCAGACCATGCGCATCAAGGTGGTGGAAGGTGG
CCCACTGCCATTCGCCTTCGATATCCTGGCCACCAGCTTTCTGTACGGCAGCAAGACCTTCATCAATCACACCCAGGGCATCCCC
GATTTCTTCAAGCAGAGCTTCCCCGAGGGCTTCACATGGGAGCGCGTGACCACATATGAGGATGGCGGAGTGCTGACCGCCACGC
AGGATACAAGTCTGCAGGATGGCTGCCTGATCTACAACGTGAAGATCCGCGGCGTGAACTTCACCAGCAATGGCCCCGTGATGCA
GAAGAAAACCCTCGGCTGGGAAGCCTTCACCGAGACACTGTATCCAGCCGATGGCGGACTGGAAGGCCGCAATGATATGGCCCTG
AAGCTCGTTGGAGGCTCCCACCTGATTGCCAACGCCAAGACAACCTACCGCAGCAAGAAGCCCGCCAAGAACCTGAAGATGCCAG
GCGTGTACTACGTGGACTACCGCCTGGAACGCATCAAAGAGGCCAACAACGAGACATACGTGGAACAGCACGAGGTGGCCGTGGC
TCGCTATTGCGATCTGCCAAGCAAGCTGGGCCACAAGCTGAATGGCGGATCCATGAGCACCGAGAGCATGATCCGTGATGTGGAA
CTGGCCGAGGAAGCCCTGCCGAAGAAAACAGGTGGACCACAGGGAAGTCGCCGCTGCCTGTTTCTGAGCCTGTTCAGCTTCCTGA
TCGTGGCCGGTGCCACCACACTGTTCTGCCTGCTGCACTTCGGAGTGATCGGCAGCGGCAACGGCAATGGAAACGGAAATGGCAA
TGGCAACGGCAACGGAAATGGCAACGGCAATGAGCAGAAGCTGATCTCCGAGGAAGATCTCGGAAACGGAAACGGCAACGGAAGC
CTCGAGAGCAACTTCTTCGATCTGGAATGCAAGGGCATTTTCAACAAGACCATGTTCTTCCGGCTGGACCGCATCTGCGAGGATT
GCTACCAGCTGTTCCGCGAAACCAGCATCCACCGCCTGTGCAAGCAAGAGTGCTTCGGCAGCCCCTTCTTCAACGCCTGCATTGA
GGCCCTGCAACTGCACGAGGAAATGGACAAGTACAACGAGTGGCGCGATACCCTGGGCCGCAAGTAATCTAGAAA
```

Stop *Xba*I

**Figure S8: Design of a membrane tethered-saITP construct.**

**A**, Amino-acid sequence of membrane-tethered saITP. This construct comprises an N-terminal mTagBFP2 fluorophore, a transmembrane domain, an extracellular linker, and the saITP sequence. No additional cysteine residues were included in the extracellular spacer to preserve proper disulfide bridge formation in saITP. **B**, The DNA sequence of the codon-optimized membrane-tethered saITP construct. The coding sequence was synthesized as a cassette comprising an *Eco*RI site, a *Drosophila* Kozak sequence, a *Drosophila*-optimized tethered-saITP coding region, and an *Xba*I site.

**Table S1: rGC proteins used for phylogenetic analysis.**

| Species | Protein name / Gene ID | GenBank accession number |
|---|---|---|
| *Drosophila melanogaster* | CG3216 | NP_726013.2 |
| | CG31183 | NP_001287342.1 |
| | CG33958 | NP_001033958.1 |
| | CG34357 | NP_001189166.1 |
| | EHR | NP_648653.2 |
| | Gyc32E | NP_001097148.1 |
| | Gyc76C* | NP_001163473.1 |
| *Aedes aegypti* | LOC5565598 | XP_021693787.1 |
| | LOC5565611 | XP_021693784.1 |
| | LOC5566347 | XP_021700770.1 |
| | LOC5570561* | XP_021708902.1 |
| | LOC5570582 | XP_021704722.1 |
| | LOC5574680 | XP_021701637.1 |
| | LOC5577087 | XP_021706999.1 |
| | LOC5579939 | XP_001652228.1 |
| *Bombyx mori* | BmGC-I | XP_037867807.1 |
| | LOC101736434* | XP_021202770.2 |
| | LOC101739209 | XP_021206683.1 |
| | LOC101740270 | XP_004924483.2 |
| | LOC101746180 | XP_037869394.1 |
| | LOC110386620 | XP_037872650.2 |
| *Tribolium castaneum* | Gyc32E | XP_015837765.1 |
| | LOC660007* | XP_008198489.1 |
| | LOC660192 | XP_015838385.2 |
| | LOC661745 | XP_015835582.1 |
| | LOC664507 | XP_015837238.1 |
| | LOC658966 | XP_015839153.1 |
| | LOC100141543 | XP_015836546.1 |
| *Apis mellifera* | LOC409422 | XP_026295331.1 |
| | LOC550964 | XP_016766759.2 |
| | LOC726069 | XP_016772021.1 |
| | LOC724603* | XP_016766757.2 |
| | LOC100576683 | XP_026296763.1 |
| *Pediculus humanus corporis* | Phum_PHUM099980* | XP_002423970.1 |
| | Phum_PHUM100000 | XP_002423972.1 |
| | Phum_PHUM360550 | XP_002428033.1 |
| | Phum_PHUM574110 | XP_002432156.1 |
| | Phum_PHUM591960 | XP_002432558.1 |
| *Daphnia pulex* | LOC124188298 | XP_046436786.1 |
| | LOC124188825* | XP_046437660.1 |
| | LOC124196298 | XP_046447193.1 |
| | LOC124201173 | XP_046453613.1 |
| | LOC124205465 | XP_046458866.1 |
| *Homo sapiens* | GUCY2C | NP_004954.2 |
| | GUCY2D | NP_000171.1 |

| | | |
|---|---|---|
| | GUCY2F | NP_001513.2 |
| | NPR1 | NP_000897.3 |
| | NPR2 | NP_003986.2 |
| *Mus musculus* | Gucy2c | NP_001120790.1 |
| | Gucy2d | NP_001124165.1 |
| | Gucy2e | NP_032218.2 |
| | Gucy2f | NP_001007577.1 |
| | Gucy2g | NP_001074545.1 |
| | Npr1 | NP_032753.5 |
| | Npr2 | NP_776149.1 |
| *Danio rerio* | gc2 | XP_021333059.1 |
| | gucy2ca | XP_073799927.1 |
| | gucy2cb | XP_696628.5 |
| | gucy2d | XP_005165318.1 |
| | gucy2f | NP_571939.2 |
| | gucy2g | XP_009305067.2 |
| | npr1a | NP_001038402.1 |
| | npr1b | XP_009290669.2 |
| | npr2 | XP_009300100.2 |
| *Drosophila melanogaster* sGC (Outgroup) | Gyc88E | NP_001036711.1 |
| | Gyc89Da | NP_001036719.1 |
| | Gyc89Db | NP_650551.1 |
| | Gycα99B | NP_477088.2 |
| | Gycβ100B | NP_524603.2 |

* (Gyc76C subfamily clade proteins)
